# Tectonic setting shapes microbial biosynthetic potential across global geothermal environments

**DOI:** 10.1101/2025.09.14.675129

**Authors:** Ana Clara Pelliciari Silva, Benoit de Pins, Francesco Montemagno, Flavia Migliaccio, Martina Cascone, Deborah Bastoni, Bernardo Barosa, Matteo Selci, Costantino Vetriani, Agostina Chiodi, Federico A. Vignale, Maria Garcia Alai, Alberto Vitale Brovarone, Gerdhard L. Jessen, Jenny M. Blamey, CoEvolve Project consortium, J. Maarten de Moor, Karen G. Lloyd, Peter Barry, Donato Giovannelli

## Abstract

Microbial communities in geothermal environments constitute an underexplored reservoir of biosynthetic gene clusters with significant biotechnological potential. Here, we investigate the secondary metabolite potential of 219 microbial communities across marine and continental geothermal field sites, encompassing broad environmental gradients in temperature (4.7 °C to 93.5 °C), pH (0.85 to 10.3), and tectonic setting, including volcanic arcs, backarcs, divergent margins at on-axis mid-ocean ridges, post-subduction extensional arcs, and paleo-convergent intraplate plume systems. We identified 9,019 putative new biosynthetic gene cluster families, mostly lacking similarity to known biosynthetic gene clusters. Volcanic arc systems consistently exhibit the highest diversity of biosynthetic repertoires, whereas intraplate plume systems showed a greater representation of terpene-associated gene cluster families. In contrast, divergent margin systems were primarily characterized by nonribosomal peptide synthetases and ribosomally synthesized and post-translationally modified peptides pathways, together accounting for a large fraction of their predicted biosynthetic diversity. These findings suggest that tectonic context could be associated with large-scale patterns in microbial biosynthetic potential and provide a geobiological framework for guiding natural product discovery in geothermal ecosystems.

## Introduction

Microorganisms produce a vast repertoire of specialized secondary metabolites that mediate interactions with neighbouring cells and the surrounding environment and have profoundly impacted medicine and biotechnology^1,2^. The genes encoding them are typically not required for the organism’s survival and growth, but confer critical adaptive advantages, for example, by inhibiting competitors, scavenging nutrients, detoxifying metals or conferring stress tolerance, thereby enhancing the overall fitness of the producing organism^2^. Notably, natural products derived from microorganisms remain indispensable as medicinal drug prospects; indeed, ∼70 % of clinically used anti-infectives originate from naturally occurring microbial metabolites^3^. Extreme environments are also increasingly recognized as reservoirs of new secondary metabolites and potentially novel chemistries^4,5^. Across deserts^1^, deep-sea systems^6^, polar soils^4^, and hypersaline habitats^5^, genome mining and cultivation studies have revealed extensive unexplored biosynthetic diversity^1^.

Biosynthetic gene clusters (BGCs) encode the enzymatic pathways that assemble specialized metabolites^8^. Key classes of BGCs include those for nonribosomal peptide synthetases (NRPS), which encode siderophores and other metallophores^9^; polyketide synthases (PKS), that generate a wide spectrum of bioactive molecules, including many clinically important antibiotics^2,9^; ribosomally synthesized and post-translationally modified peptides (RiPPs), encompassing bacteriocins, lantibiotics, and archaeocins, which frequently act as antimicrobials that shape microbial communities, particularly in dense biofilms^10–13^ and; terpene synthase clusters that produce structurally diverse metabolites involved in signaling, membrane stabilization, and oxidative stress protection^14^. Prior work has highlighted geothermal extremophiles as an untapped source of natural products^15,16^, with environmental stressors often correlating with enrichment of specific BGC classes in the microbiome: for example, chronic metal exposure favors NRPS-linked metallophores that detoxify metals^16^, whereas extremely acidic, heat-stressed systems dominated by few chemolithotrophs often show a dearth of large NRPS/PKS clusters^17,18^. These patterns suggest that BGC repertoires reflect the selective pressures of their environment. It is important to notice, however, that the geochemical landscapes in which microbial communities adapt to are not random but are firmly rooted in the geological settings from which they arise^19^. The mineral content, redox potential, temperature and ionic composition of each environment are shaped by underlying tectonic and geological processes, which in turn influence microbial community structure and function^19,20^.

There is now compelling evidence that the microbiome of extreme environments has the potential to yield a wealth of previously unknown natural products^4^. For instance, a recent review paper documented 186 new secondary metabolite structures isolated from 129 extremophilic strains collected between 2010 and 2018^1^. Among these discoveries are novel antimicrobial and cytotoxic compounds from desert actinomycetes, including 46 new molecules identified from Atacama Desert isolates alone^5^. Likewise, organisms from other extreme biomes, such as Antarctic soil communities and deep-sea sediments, have revealed extraordinary biosynthetic richness, with little overlap to known natural products^7^. Even archaeal lineages, with secondary metabolism historically underexplored, have been found to harbor BGCs in extreme settings: for example, *Heimdallarchaeota* and *Lokiarchaeota* from hypersaline microbial mats were recently shown to possess putatively novel BGCs, expanding the phylogenetic range of natural product producers^4^.

In spite of increasing evidence suggesting the high potential of extreme environments in harboring new biosynthetic pathways and metabolites, large scale BGCs surveys across diverse ecosystems are lacking. Advances in genomics and metagenomics are now enabling the systematic exploration of BGC potential in these inaccessible ecosystems^21^. Many extremophiles remain uncultivated, but culture-independent genome mining approaches, including high-throughput sequencing of environmental DNA and reconstruction of metagenome-assembled genomes (MAGs), allow researchers to recover biosynthetic genes directly from environmental samples^22^. Using these approaches, recent works explored the biosynthetic potential of microbiomes on a global scale, across terrestrial and marine environments^23, 22^ and in more localized settings such as the extreme microbial mats of Shark Bay^2^, Antarctic soils^24^, glacier-fed stream biofilms^25^, or the Tibetan Plateau extreme aquatic microbiomes^26^. Although geothermal environments have played a key role over the last decades as sources of bioactive compounds and enzymes used in industry^27–29^, we lack systematic investigations of the biosynthetic potential of geothermal systems and how environmental factors shape the diversity and distribution of predicted BGCs.

Here, we address this knowledge gap by investigating the biosynthetic potential of global geothermal extreme environments. We combine comprehensive analyses of 219 microbial communities from marine and continental geothermal sites to map the diversity and distribution of predicted BGCs from distinct gene cluster families (GCFs) across diverse tectonic settings. Our results show that variation in the tectonics and geochemical context might be associated with distinct patterns in composition and diversity of predicted microbial natural products. This work lays a foundation for future bioprospecting efforts and expands our understanding of how Earth’s subsurface could influence chemical innovation in the microbial world.

## Results

We analyzed 219 microbial communities from geothermal features across ten major geothermal regions using a combination of metagenomic sequencing and geochemical measurements (Figure 1). These include environments in present-day volcanic arcs (*e.g.*, the South American Central Volcanic Zone (SA-CVZ), Central America Volcanic Arc (CAVA), and Aeolian Arc Volcanic Province (AAVP)), adjacent backarc regions (*e.g.,* part of the SA-CVZ), and transitional post-subduction extensional arcs (Campania, southern Italy and the Tuscan-Latium Volcanic Province, (TLVP) in central Italy); on-axis mid-ocean ridge settings (*e.g.*, Reykjanes Volcanic Belt, (RVB), deep-sea vents in the Guaymas Basin and the East Pacific Rise, (EPR)); and intracontinental plume systems in paleoaccretionary environments in the South Khangai Volcanic Province (SKVP) (Figure 1).

**Figure 1.**
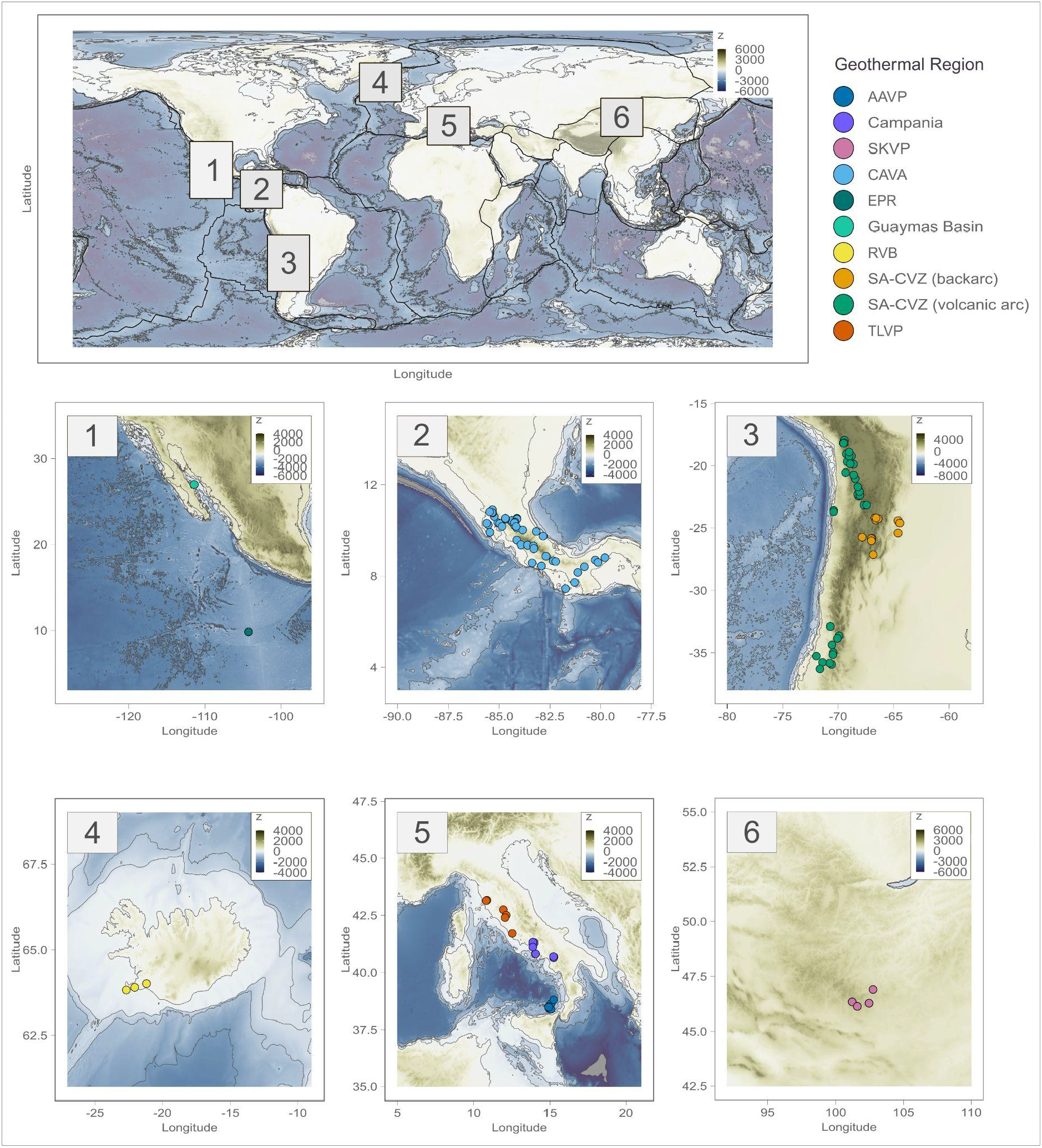
Geographic distribution of sampled geothermal extreme environments included in this study. Locations span diverse geothermal regions in the Guaymas basin and the East Pacific Rise (EPR) (1), the Central American Volcanic Arc (CAVA) (2), the South American Central Volcanic Zone (SA-CVZ) (3), Reykjanes Volcanic Belt (RVB) (4), the Tuscan-Latium Volcanic Province (TLVP), the Campania region, and the Aeolian Arc Volcanic Province (AAVP) (5), and the South Khangai Volcanic Province (SKVP) (6).

Biosynthetic potential was characterised using presence-absence profiles of gene cluster families (GCFs). Partial redundancy analysis (RDA) revealed distinct patterns linked to tectonic context and geochemical regimes (overall model: F = 1.69, p < 0.001; Figure 2). Volcanic arc samples preferentially occupy distinct clusters from those of backarc, post-subduction extensional volcanics, and on-axis mid-oceanic ridge (MOR) spreading systems. The observed clusters differed significantly in biosynthetic composition (PERMANOVA on Hellinger-Euclidean distances R² = 0.039, p < 0.001; ANOSIM: R = 0.26, n = 219, adj.p < 0.001), showing that the geothermal systems located in diverse tectonic systems might be characterised by distinct combinations of putative GCFs, as well their overall predicted biosynthetic potential.

**Figure 2.**
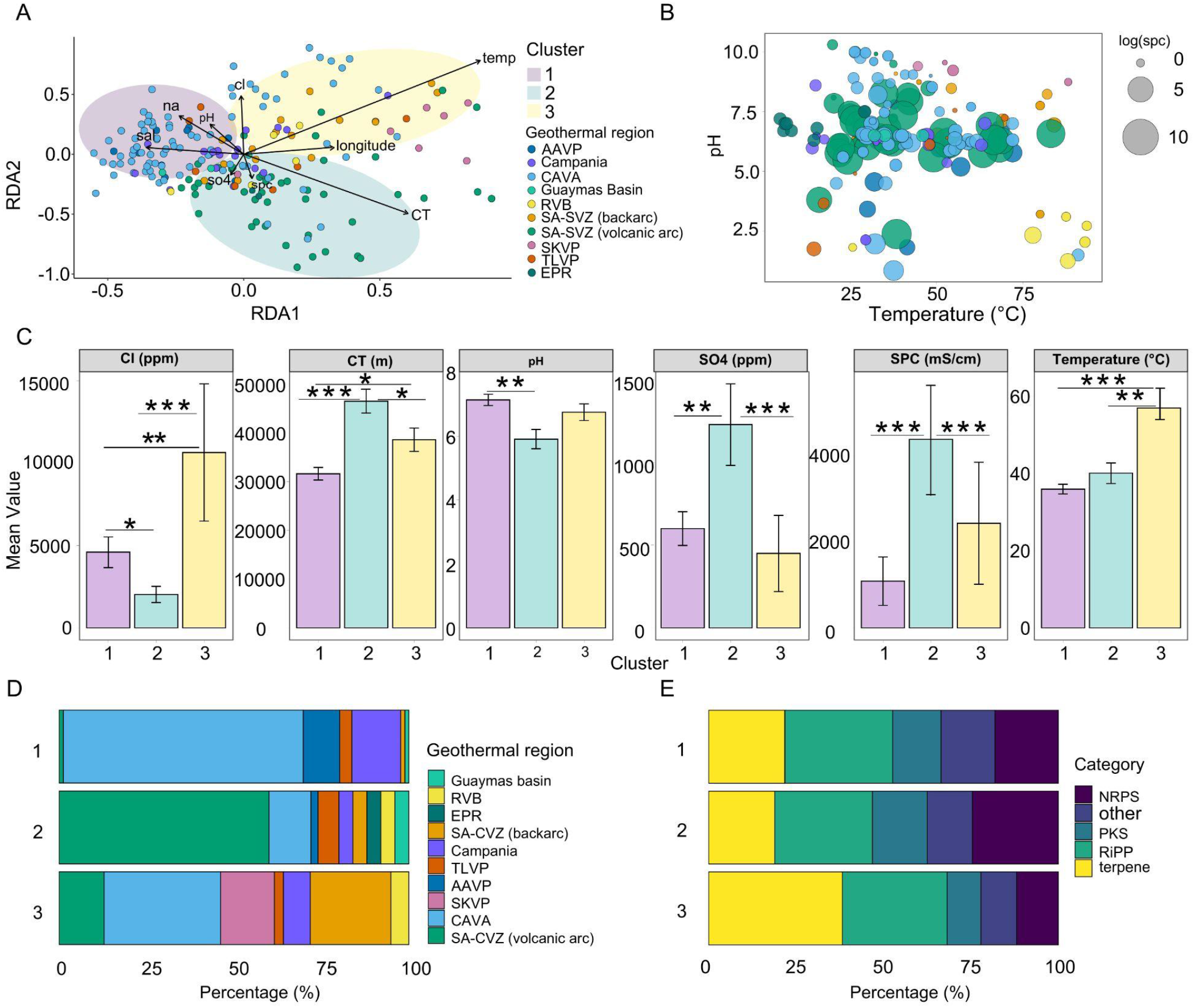
Biosynthetic gene cluster family (GCF) composition across geothermal sites. **(a)** Partial RedundancyAnalysis (RDA) of GCF composition across geothermal metagenomes. Points are colored by geothermal regions of origin: CAVA (Central American Volcanic Arc), SA-CVZ (South American Central Volcanic Zone), RVB (Reykjanes Volcanic Belt), TLVP (Tuscan-Latium Volcanic Province), AAVP (Aerolian Arc Volcanic Province), Campania region, SKVP (South Khangai Volcanic Province), the Guaymas basin, and the EPR (East Pacific Rise). Arrows represent environmental variables, with direction and length proportional to their correlation with ordination axes (*e.g.*, pH, temperature, salinity, conductivity (spc), crustal thickness (CT), elevation, Cl^−^ content, SO_4_ content, longitude). Colored convex hulls illustrate the spatial distribution of three k-means clusters. **(b)** pH as a function of temperature scatter plot. Size of samples are correlated with their log conductivity values (spc), and coloured by geothermal region of origin. **(c)** Environmental variables by cluster. Mean ± standard error (SE) values are shown for six key environmental parameters across microbial community clusters (1–3). Colors represent cluster identity: pale yellow (Cluster 1), pale teal (Cluster 2), and pale purple (Cluster 3). Statistical differences were tested using the Kruskal-Wallis test followed by Dunn’s post-hoc comparisons with Benjamini-Hochberg correction. Asterisks indicate significant pairwise differences between clusters (*p < 0.05, **p < 0.01, ***p < 0.001). **(d)** Geothermal region origin across tectonic clusters. Stacked horizontal bar plots show the proportion of each geothermal region represented within each k-means-derived cluster. **(e)** Relative distribution of major predicted GCF classes across sample clusters, reflecting presence/absence of biosynthetic profiles.

Three major BGC clusters were identified along the main axis of the ordination, each corresponding to different geochemical and tectonic regimes (Figure 2a-d). Samples in cluster 1 and 2, are both dominated by volcanic arc systems (>80 % of samples), but differed in environmental associations: Cluster 1 is associated with higher pH, whereas Cluster 2 is associated with higher electrical conductivity (spc), and crustal thickness (CT) values (Figure 2b-c). In contrast, Cluster 3 includes a stronger presence of geothermal regions associated with backarc margins and intracontinental plume settings, aligning closely with high-temperature environments, suggesting it includes predominantly sites that could host thermophilic microbes.

We next examined the biosynthetic composition across clusters. NRPS-associated GCFs are most frequent in Cluster 2 (26 %), followed by Cluster 1 (19 %), and Cluster 3 (12 %). In contrast, terpene-associated GCFs have higher representation in Cluster 3 (39 %), followed by Cluster 1 (21 %) and Cluster 2 (18 %). RiPP-associated GCFs are broadly distributed across all clusters (28-31 %), whereas PKS-associated GCFs are less prevalent overall (9-15 %). The remaining GCFs, which includes hybrid and, not yet fully categorised clusters, grouped as “Other,” comprise a smaller fraction of the putative biosynthetic repertoire (10-18 % across clusters), with Cluster 1 showing the highest representation (18%), followed by Cluster 2 (15%) and Cluster 3 (10%) (Figure 2e).

Among the environmental variables tested, temperature (range: 4.7 °C to 93.5 °C), pH (range: 0.85 to 10.3), and crustal thickness (range: 7.44 km to 78.26 km) show consistent associations with variation in predicted biosynthetic GCF profiles. Marginal PERMANOVA analyses reveal that temperature explains the largest proportion of variance (R² = 0.012, F = 2.5, adj.p < 0.001; Supplementary Figure 1 top), followed by pH (R² = 0.010, F = 2.07, adj.p < 0.001; Supplementary Figure 1 middle ) and crustal thickness (CT; R² = 0.007, F = 1.33, adj. p < 0.01). Although each variable individually accounted for a modest fraction of total variance, their combined effects define a multidimensional environmental gradient that might have an influence on biosynthetic repertoires across geothermal systems (Supplementary Figure 1). Notably, these variables can covary across tectonic environments, which can reflect linked geological processes that influence fluid chemistry and thermal regimes. When analysing samples in active convergent margins, volcanic arc geothermal systems exhibit greater putative GCF diversity in comparison to backarc and post-subduction extensional arcs (volcanic arc: 2.18 ± 0.69; backarc: 1.85 ± 0.57; post-subduction extensional arcs: 1.88 ± 0.41; Kruskal–Wallis χ² = 17.35, adj.p = 1.7 × 10⁻^4^, Supplementary Figure 2).

Microbial communities from volcanic arc springs reveal significantly higher predicted NRPS presence compared with all other sites (Figure 3), whereas the putative metagenomic GCF profile of backarc sites shows elevated terpene presence compared to other active plate tectonic sites (Figure 3). Geochemical characterization of representative geothermal systems reveals distinct environmental conditions across tectonic settings (Table 1, Supplementary Tables 1–4, Supplementary Figure 3). Volcanic arc systems (*e*.*g*, part of the SA-CVZ and AAVP) are characterized by elevated concentrations of dissolved trace metals, and backarc geothermal systems exhibit CO₂-dominated gas compositions (62–99 mol% CO₂). Geothermal systems associated with on-axis ridges such as the RVB also contained elevated concentrations of trace metals, including Fe, Mn, and Zn. These geochemical differences provide environmental context for the unique biosynthetic profiles observed across tectonic settings. Interestingly, our analysis of the SKVP geothermal sites (the only sampling location present within a paleoaccretionary plume tectonic setting rather than active plate tectonics) reveals the most elevated presence of putative terpene-associated GCFs (47 %), a value nearly double that observed in tectonically active volcanic arcs, where terpene contributions never exceed 25 % (Figure 3, Supplementary Figure 9).

**Figure 3.**
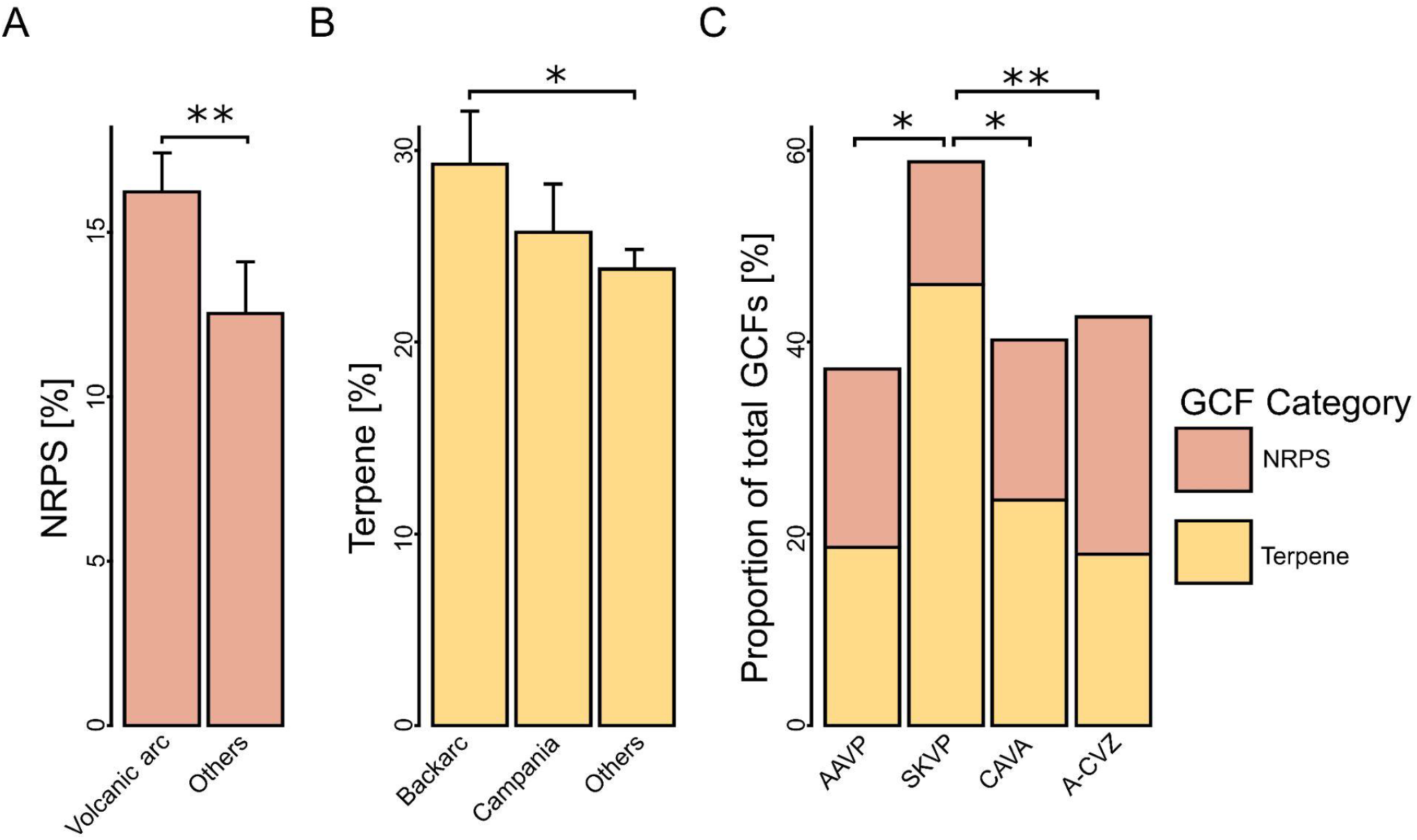
Relative representation of NRPS- and terpene-associated gene cluster families across tectonic settings and geothermal regions. Per-sample relative representation was calculated as the number of GCFs assigned to the indicated biosynthetic class divided by the total number of GCFs detected in each sample (%). **(a)** Relative representation of NRPS-associated GCFs in volcanic arc samples (n = 125) compared with samples from other active plate-tectonic settings (n = 87). Differences were assessed using a two-sided Wilcoxon rank-sum (Mann–Whitney U) test. **(b)** Relative representation of terpene-associated GCFs in backarc (n = 34), Campania (n = 17), and other active plate-tectonic settings (n = 161). Overall differences among groups were assessed using a Kruskal–Wallis test, followed by Dunn’s post hoc test with Holm correction for multiple comparisons. **(c)** Relative representation of terpene- and NRPS-associated GCFs across selected geothermal regions: AAVP (Aeolian Arc Volcanic Province; n = 10), SKVP (n = 7), CAVA (Central American Volcanic Arc; n = 88), and SA-CVZ (South American Central Volcanic Zone; n = 46). Differences among geothermal regions were assessed using a Kruskal–Wallis test followed by Dunn’s post hoc test with Holm correction for multiple comparisons. Significant pairwise comparisons are indicated by p <0.05 (*), p < 0.01 (**).

**Table 1.** Coordinates and basic parameters of selected sampled sites.

| Site name | Geothermal region | Latitude (°N) | Longitude (°E) | Temperature (°C) | pH | Cl (ppm) | Na (ppm) | K (ppm) | Mg (ppm) | Ca (ppm) | Mn (ppb) | Fe (ppb) | Zn (ppb) | CO<sub>2</sub> (mmol mol<sup>-1</sup>) |
| --- | --- | --- | --- | --- | --- | --- | --- | --- | --- | --- | --- | --- | --- | --- |
| Acque Calde | AAVP | 38.42 | 14.96 | 30.0 | 3.4 | 5009.7 | 3644.0 | 172.0 | 585.9 | 570.2 | 2082.4 | 54138.5 | 44.3 | 860.3 |
| Bottaro | AAVP | 38.38 | 15.06 | 68.4 | 6.7 | 20664.4 | 12185.9 | 415.7 | 2147.8 | 965.7 | 0.0 | 0.0 | 0.0 |  |
| Black Point White | AAVP | 38.38 | 15.06 | 32.0 | 4.6 | 21056.2 | 11735.1 | 459.3 | 1923.1 | 1171.1 |  | 0.0 | 0.0 |  |
| Bagno Secco Sorgente Rossa | AAVP | 38.49 | 14.91 | 46.2 | 6.6 | 173.8 | 3637.4 | 42.2 | 91.0 | 191.8 |  |  |  |  |
| Casa Fulco | AAVP | 38.80 | 15.24 | 40.5 | 6.3 | 6077.7 | 3567.2 | 43.2 | 1101.4 | 999.4 | 1730.7 | 81.4 | 123.9 | 963.2 |
| Fangaia Vulcano | AAVP | 38.42 | 14.96 | 41.5 | 1.8 | 2204.9 | 1383.2 | 160.0 | 200.8 | 300.7 | 2060.3 | 93113.9 | 0.0 | 716.0 |
| Geyser Vulcano | AAVP | 38.42 | 14.96 | 72.0 | 5.8 | 21850.8 | 12134.3 | 424.4 | 2082.2 | 906.5 |  |  |  |  |
| Hotel Oasi | AAVP | 38.64 | 15.08 | 50.8 | 6.0 | 14782.4 | 4252.8 | 313.6 | 1070.9 | 743.9 | 6560.7 | 0.0 | 0.0 | 981.1 |
| Levante Bay | AAVP | 38.42 | 14.96 | 57.0 | 5.19 | 20338.3 | 11196.9 | 470.9 | 1820.4 | 872.7 | 702.3 | 180.6 | 0.0 | 982.8 |
| Terme San Calogero Stufa | AAVP | 38.48 | 14.91 | 37.8 | 7.8 | 218.8 | 368.2 | 33.9 | 631.5 | 588.5 | 106.2 | 0.0 | 0.0 | 711.2 |
| Villa Levante | AAVP | 38.41 | 14.96 | 64.4 | 3.4 | 27660.0<br>2 | 9763.1 | 889.0 | 1978.3 | 778.9 |  |  |  |  |
| Gunnuhver | RVB | 63.82 | -22.68 | 77.9 | 2.3 | 2601.03 | 1599.0 | 172.7 | 56.7 | 189.0 | 1531.9 | 91035.7 | 560.8 |  |
| Gunnuhver | RVB | 63.82 | -22.68 | 88.0 | 1.3 | 1398.8 | 909.0 | 98.8 | 34.9 | 93.7 | 1014.1 | 68824.5 | 1428.3 |  |
| Hveragerdi | RVB | 64.01 | -21.18 | 93.5 | 2.7 | 1.5 | 8.7 | 1.5 | 44.5 | 71.8 | 2984.2 | 2716.6 | 8607.4 |  |
| Hveragerdi | RVB | 64.01 | -21.18 | 25.7 | 1.8 | 2.4 | 19.7 | 7.6 | 68.4 | 107.2 | 3974.5 | 39773.4 | 555.0 |  |
| Krysuvik | RVB | 63.89 | -22.06 | 87.5 | 3.1 | 6.9 | 6.8 | 0.93 | 9.4 | 12.1 | 393.6 | 24729.0 | 1113.3 |  |
| Krysuvik | RVB | 63.90 | -22.06 | 93 | 2.1 | 3.6 | 5.3 | 0.8 | 6.0 | 13.2 | 205.4 | 21148.3 | 944.7 |  |
| Las Cuevas | SA-CVZ | -18.17 | -69.43 | 33.7 | 6.4 | 368.8 | 316.9 | 6.6 | 8.7 | 65.2 | 3.2 | 11.9 | 581.6 |  |
| Termales Chusmiza | SA-CVZ | -19.68 | -69.18 | 41.7 | 8.2 | 63.2 | 127.0 | 4.1 | 0.1 | 27.1 | 9.1 | 0.2 | 1021.4 |  |
| Termales Enquelga | SA-CVZ | -19.23 | -68.79 | 30.1 | 6.0 | 177.3 | 172.0 | 40.8 | 63.6 | 140.5 | 5.8 | 23.4 | 378.04 |  |
| El Tatio | SA-CVZ | -22.33 | -68.01 | 86.8 | 6.3 | 11255.5 | 3874.9 | 514.6 | 0.0 | 226.4 | 420.8 | 13.0 | 354.6 | 6.0 |
| Geyser Blanco | SA-CVZ | -22.36 | -68.02 | 83.1 | 6.6 | 2146.0 | 987.5 | 90.6 | 12.3 | 80.6 | 350.3 | 37.1 | 836.7 | 495.4 |
| Irruputuncu acid spring | SA-CVZ | -20.73 | -68.59 | 38.4 | 2.4 | 1995.7 | 747.3 | 136.8 | 458.8 | 221.4 | 33041.5 | 38515.7 | 1416.3 |  |
| Termales Jurasi | SA-CVZ | -18.21 | -69.51 | 65.4 | 6.9 | 266.5 | 275.3 | 9.9 | 0.9 | 59.7 | 617.7 | 56.3 | 2.9 |  |
| Lagu Amarilla | SA-CVZ | -19.06 | -69.25 | 35.5 | 6.8 | 5497.6 | 2043.9 | 268.8 | 35.9 | 660.2 | 2435.1 | 328.2 | 390.2 |  |
| La Chimba | SA-CVZ | -23.56 | -70.40 | 40.3 | 7.3 | 5716.6 | 2602.6 | 80.0 | 140.3 | 463.2 | 17.9 | 0.05 | 62.05 |  |
| Lagu Negra | SA-CVZ | -23.15 | -67.42 | 52.3 | 6.6 | 17248.4 | 5345.8 | 130.8 | 127.7 | 1335.2 | 6193.9 | 827.3 | 647.1 | 721.1 |
| Lagu Paricota | SA-CVZ | -19.23 | -69.01 | 24.9 | 7.5 | 64.9 | 42.6 | 8.6 | 7.0 | 22.9 | 16.6 | 53.0 | 1063.9 |  |
| Lagu Roja | SA-CVZ | -19.85 | -68.91 | 24.3 | 5.5 | 6130.9 | 1929.9 | 254.2 | 35.0 | 624.7 | 1630.5 | 27.4 | 402.3 |  |
| Lagu Verde | SA-CVZ | -19.06 | -69.25 | 49.1 | 5.6 | 4353.1 | 1686.4 | 220.4 | 30.1 | 558.6 | 2249.4 | 183.2 | 406.3 | 963.4 |
| Ojo de Ascotán vertiente 11 | SA-CVZ | -21.69 | -68.22 | 27.5 | 7.0 | 1056.2 | 332.7 | 31.9 | 37.6 | 142.4 | 5.4 | 43.0 | 493.9 | 871.7 |
| Parajaya | SA-CVZ | -19.12 | -68.91 | 29.6 | 6.7 | 146.0 | 114.0 | 18.5 | 53.6 | 0.0 | 2.9 | 5.2 | 350.3 | 174.6 |
| Baño Polio Quieres | SA-CVZ | -18.91 | -69.00 | 63.4 | 5.8 | 2187.3 | 1099.4 | 191.6 | 46.7 | 197.7 | 2705.7 | 544.0 | 694.0 | 930.3 |
| Putre | SA-CVZ | -18.20 | -69.54 | 35.8 | 6.3 | 5077.8 | 1948.4 | 220.0 | 31.0 | 349.0 | 1575.6 | 4528.7 | 782.9 | 982.4 |
| Quebrada Negra | SA-CVZ | -23.70 | -70.41 | 26.4 | 7.1 | 16229.0 | 4511.2 | 72.7 | 241.9 | 1539.7 | 6.4 | 41.0 | 509.3 |  |
| Rio Salado | SA-CVZ | -22.28 | -68.23 | 22.24 | 6.0 | 1969.0 | 922.2 | 71.2 | 77.7 | 250.6 | 6409.0 | 30.7 | 715.5 | 937.6 |
| Santa Rosita | SA-CVZ | -20.54 | -69.33 | 34.8 | 8.3 | 1195.9 | 570.8 | 5.0 | 1.3 | 296.3 | 8.7 | 97.2 | 1319.9 |  |
| Termales Lirima | SA-CVZ | -19.85 | -68.91 | 68.7 | 6.3 | 334.7 | 291.8 | 32.2 | 2.1 | 42.4 | 168.7 | 91.2 | 864.3 | 484.3 |
| Termales Mamiña | SA-CVZ | -20.07 | -69.21 | 53.9 | 8.8 | 131.5 | 84.5 | 1.9 | 0.0 | 19.6 | 3.5 | 11.8 | 646.7 |  |
| Termal Tana | SA-CVZ | -19.11 | -69.14 | 66.6 | 6.3 | 4050.6 | 1488.7 | 183.2 | 15.0 | 249.9 | 2211.6 | 2540.4 | 709.2 |  |
| Vertiente 10 Ascotán | SA-CVZ | -21.61 | -68.25 | 22.2 | 7.2 | 2551.2 | 816.04 | 69.0 | 72.2 | 239.3 | 1.1 | 39.2 | 242.6 |  |
| Antuco | SA-CVZ (backarc) | -24.18 | -66.67 | 27.8 | 6.3 | 9367.5 | 5162.9 | 603.8 | 154.9 | 414.1 | 289.5 | 1658.1 | 0.0 | 989.6 |
| Botijuela | SA-CVZ (backarc) | -25.74 | -67.82 | 40.0 | 6.4 | 1590.8 | 1508.0 | 84.1 | 21.9 | 86.8 | 183.1 | 228.9 | 0.0 | 979.5 |
| Galán Aguas Calientes | SA-CVZ (backarc) | -25.83 | -66.92 | 67.0 | 6.7 | 647.3 | 533.3 | 10.3 | 1.1 | 16.1 | 98.4 | 33.1 | 0.0 | 952.9 |
| Galán Fumaroles (El Diablo) | SA-CVZ (backarc) | -25.86 | -66.99 | 80.0 | 7.8 | 14.7 | 12.4 | 2.9 | 1.0 | 3.7 | 71.5 | 585.7 | 0.0 | 882.6 |
| Galán Fumaroles (El Diablo) | SA-CVZ (backarc) | -25.86 | -66.99 | 80.0 | 3.2 | 50.8 | 23.9 | 10.4 | 4.3 | 21.2 | 406.8 | 17292.8 | 6.0 | 910.0 |
| Incachule | SA-CVZ (backarc) | -24.28 | -66.47 | 46.9 | 6.5 | 350.3 | 424.5 | 31.5 | 4.8 | 25.9 | 117.2 | 146.0 | 0.0 | 621.2 |
| Galán La Colcha | SA-CVZ (backarc) | -26.03 | -66.99 | 84.0 | 6.9 | 1179.4 | 915.2 | 51.0 | 0.6 | 35.8 | 205.1 | 53.7 | 65.7 | 979.4 |
| Pastos Grandes | SA-CVZ (backarc) | -24.36 | -66.57 | 44.9 | 8.8 | 177.4 | 134.0 | 1.6 | 0.3 | 23.7 | 3.1 | 24.7 | 0.0 | 911.4 |
| Pompeya | SA-CVZ (backarc) | -24.25 | -66.36 | 50.3 | 6.5 | 1129.1 | 1115.5 | 112.4 | 25.8 | 45.6 | 69.3 | 298.4 | 0.0 | 978.9 |
| El Galpón | SA-CVZ (backarc) | -24.41 | -64.59 | 54.3 | 8.5 | 141.0 | 263.5 | 2.9 | 0.2 | 5.3 | 42.2 | 1552.1 | 17.7 | 957.7 |
| Rosario de la Frontera | SA-CVZ (backarc) | -25.41 | -64.59 | 82.0 | 8.2 | 424.0 | 404.3 | 5.5 | 0.2 | 5.6 | 8.4 | 99.2 | 0.0 | 954.5 |
| Tocomar | SA-CVZ (backarc) | -24.19 | -66.55 | 69.2 | 7.1 | 682.6 | 772.0 | 76.1 | 4.3 | 29.5 | 289.0 | 463.8 | 0.0 | 992.2 |
| Villa Vil | SA-CVZ (backarc) | -27.11 | -66.82 | 38.2 | 9.1 | 55.2 | 82.7 | 0.81 | 0.088 | 6.8 | 17.4 | 319.6 | 1.1 | 963.0 |

We next examined the phylogenetic diversity of representative predicted terpene biosynthetic core proteins recovered from the metagenomes (Figure 4). Phylogenetic placement alongside experimentally characterized terpene synthases reveals extensive diversity, with environmental sequences distributed across the tree and frequently forming clades lacking closely related reference proteins. While these analyses describe how putative biosynthetic gene cluster repertoires vary and diversify across environmental gradients and tectonic contexts, they do not address the extent to which this diversity might correspond to previously characterized biosynthetic pathways. We therefore next evaluated the novelty of predicted biosynthetic gene clusters by comparing the putative environmental BGCs to experimentally validated reference clusters. Network-based dereplication identified 9,019 GCFs across all samples (from 37,178 individual BGCs, Supplementary Table 5, Figure 5 and Supplementary Figures 5-7). A substantial fraction of these families show no network connections to known reference BGCs, indicating extensive unexplored biosynthetic diversity within geothermal microbiomes, an example can be seen in Figure 5. Analysis of the major biosynthetic classes revealed consistently high levels of novelty: 89% of terpene, 91% of NRPS, 94% of RiPP, and 97% of PKS GCFs (Supplementary Figures 5-7) were not connected to experimentally characterized biosynthetic gene clusters. To further assess how novelty varies across environmental gradients, we evaluated the minimum novelty per sample for each major biosynthetic class. Across all classes, relationships with temperature and pH are weak and generally not statistically significant (R² < 0.02; p > 0.05 in most cases, Supplementary Figure 8). In contrast, crustal thickness (CT) showed a consistent, albeit modest, negative relationship with minimum novelty for multiple classes, including terpene (R² = 0.033, p = 0.0107), NRPS (R² = 0.043, p = 0.0063), PKS (R² = 0.041, p = 0.0070), and RiPP (R² = 0.050, p = 0.0013) (Figure 7c, Supplementary Figures 5c–7c). These results indicate that while overall novelty remains high across geothermal environments, large-scale geodynamic context, as approximated by crustal thickness, may exert a consistent influence on the degree of novelty amongst predicted environmental GCFs. Notably, samples from mid-ocean ridge environments (EPR and Guaymas basin) tend to occupy the moral novel end of this gradient, whereas subduction-related volcanic arc systems (convergent margins in the SA-CVZ) are associated with predicted GCFs that are more closely related to experimentally characterized pathways. This pattern may also, in part, reflect underlying sampling biases in current reference databases, which are disproportionately composed of terrestrial and easily accessible environments.

**Figure 4.**
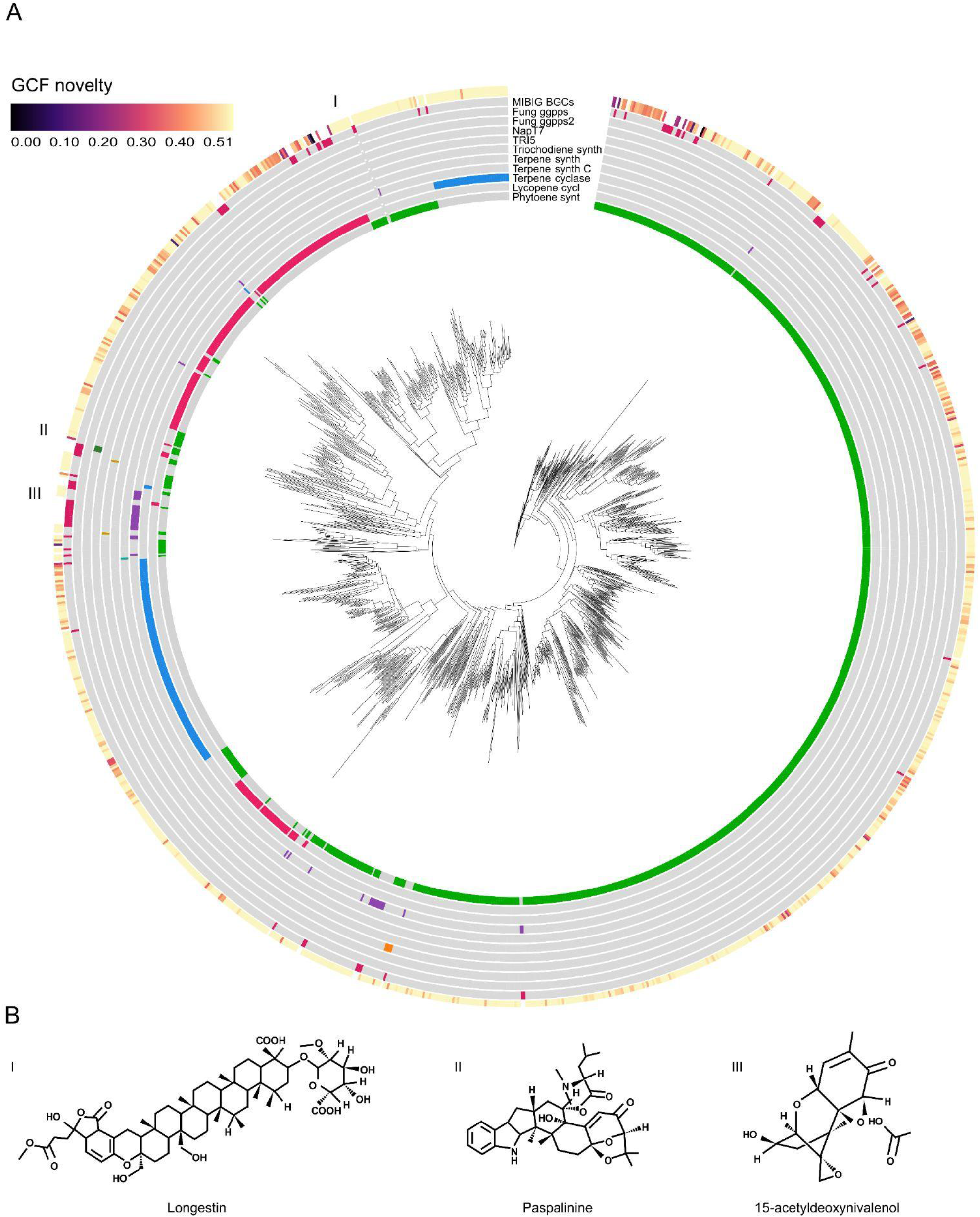
Phylogenetic diversity of representative terpene biosynthetic core enzymes and representative terpene chemistries. (a) Maximum-likelihood phylogeny of representative predicted terpene biosynthetic core proteins identified in geothermal metagenomes together with experimentally characterized reference sequences from the MIBiG (Minimum Information about a Biosynthetic Gene) database. Concentric rings indicate GCF novelty relative to MIBiG, and annotation features associated with terpene biosynthesis, including MIBiG reference clusters and functional domains identified by antiSMASH (e.g., terpene synthases, terpene cyclases, phytoene synthases). Labeled regions (I–III) highlight clades containing reference enzymes used to illustrate representative terpene chemical scaffolds. (b) Representative metabolites produced by experimentally characterized terpene biosynthetic pathways corresponding to the highlighted phylogenetic regions. These compounds (e.g., Longestin, Paspalinine, and 15-acetyldeoxynivalenol) illustrate known terpene structural diversity associated with nearby reference enzymes in the tree and provide biochemical context for the broader terpene diversity observed in geothermal microbial communities. Such compounds and related terpene scaffolds may represent candidate molecular biomarkers for microbial activity in geothermal environments.

**Figure 5.**
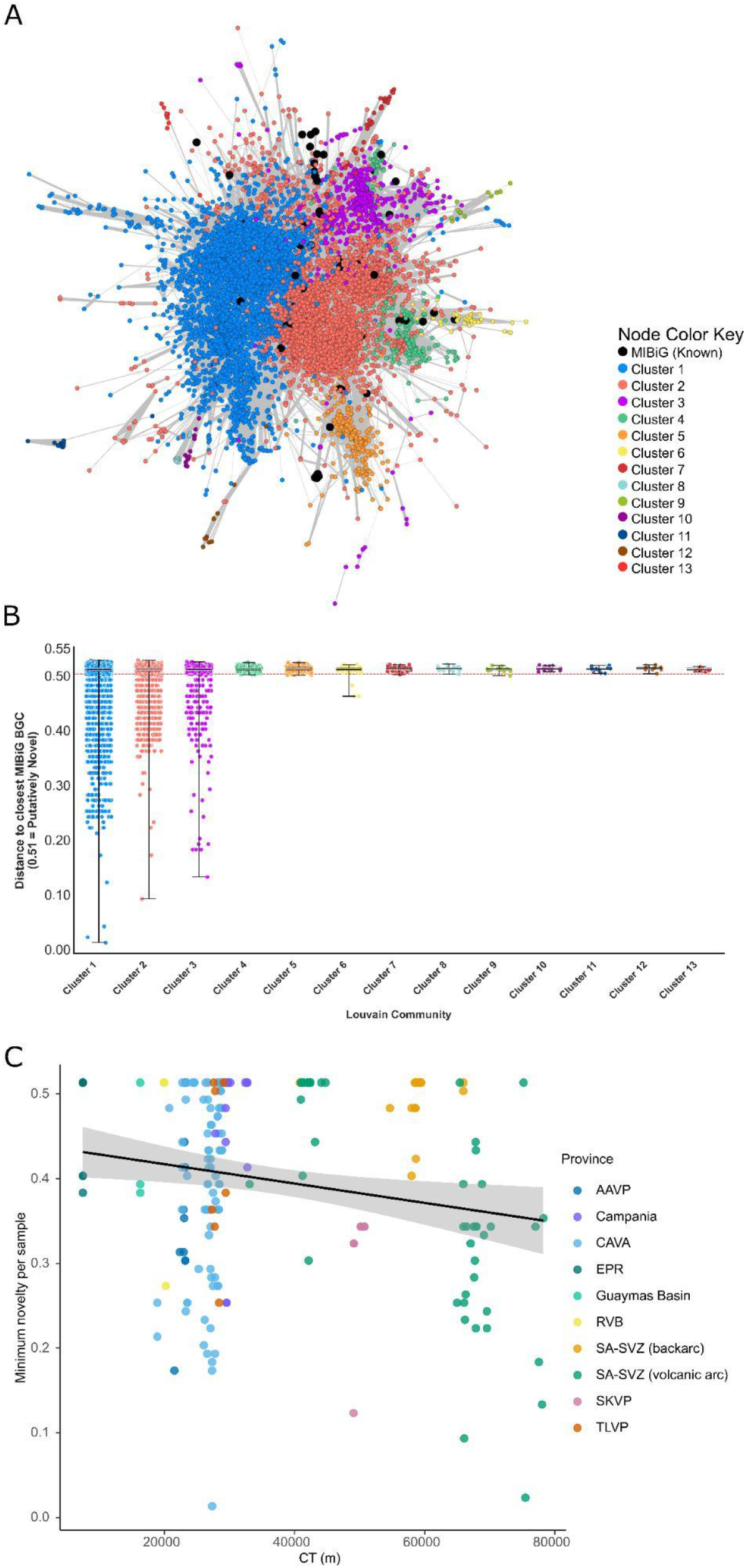
Network structure and novelty of putative terpene biosynthetic gene clusters. **(a)** Similarity network of predicted terpene biosynthetic gene clusters generated using BiG-SCAPE (Biosynthetic Gene Similarity Clustering and Prospecting Engine) at a distance cutoff of 0.5. Nodes represent BGCs (biosynthetic gene clusters) and edges indicate pairwise similarity (distance ≤ 0.5). Nodes are colored according to Louvain communities, and experimentally characterized reference clusters from the MIBiG (Minimum Information about a Biosynthetic Gene) are shown in black. Connected components correspond to predicted gene cluster families (GCFs). **(b)** Distribution of distances from each putative BGC to its closest MIBiG reference cluster, grouped by Louvain community. Each point represents a single predicted BGC, with boxplots summarizing the distribution within each community. Distances of 0.51 (above the cutoff, indicated by the dashed red line at 0.5) correspond to putative BGCs with no detectable similarity to any MIBiG cluster within the threshold. **(c)** Minimum terpene BGC novelty per sample as a function of crustal thickness (CT), colored by geothermal region. Each point represents a single sample, and the solid line indicates a linear regression with 95% confidence interval (grey shading).

## Discussion

### Geology, geochemistry and biosynthetic potential

The extreme conditions and metal bioavailability in geothermal systems of active plate tectonic sites not only select which microbes thrive^30^, but also the GCF repertoire those communities harbor. Consistent with this ecological framework, our analyses reveal that variation in biosynthetic GCF composition across global geothermal systems is associated with both environmental context and underlying taxonomic structure (Supplementary Figure 4). In particular, geothermal systems of volcanic arcs exhibit high potential for predicted microbial GCF richness (Cluster 1 and 2, Figures 2d–e, Supplementary Figure 2) and significantly greater presence of NRPS-associated predicted GCFs than other tectonic settings (Figure 3). This is exemplified by SA-CVZ volcanic arc geological context (Figure 6a) where deeply sourced springs fed by surrounding arc volcanism carry waters with high arsenic concentrations (often >1 mg/L, Supplementary Table 1) derived from arc geothermal inputs^31^, along with several metals (*e.g*., Fe, Mn, Zn) mobilized by magmatic fluids^32^. Despite the toxicity of some of these metals, the resident microbial community in the hot springs of the SA-CVZ volcanic arc is metabolically rich^33,34^. Similarly geothermal fields from other volcanic arc systems within our dataset, such as the AAVP (Figure 6b), are characterised by acidic, sulfur-rich fluids enriched in trace metals (Supplementary Table 2), consistent with previous geochemical descriptions of these systems^35^. These geological and geochemical characteristics coincide with the elevated predicted NRPS prevalence in volcanic arcs, which could suggest prolific metallophore production and metal-microbe interactions in this environment^36^.

**Figure 6.**
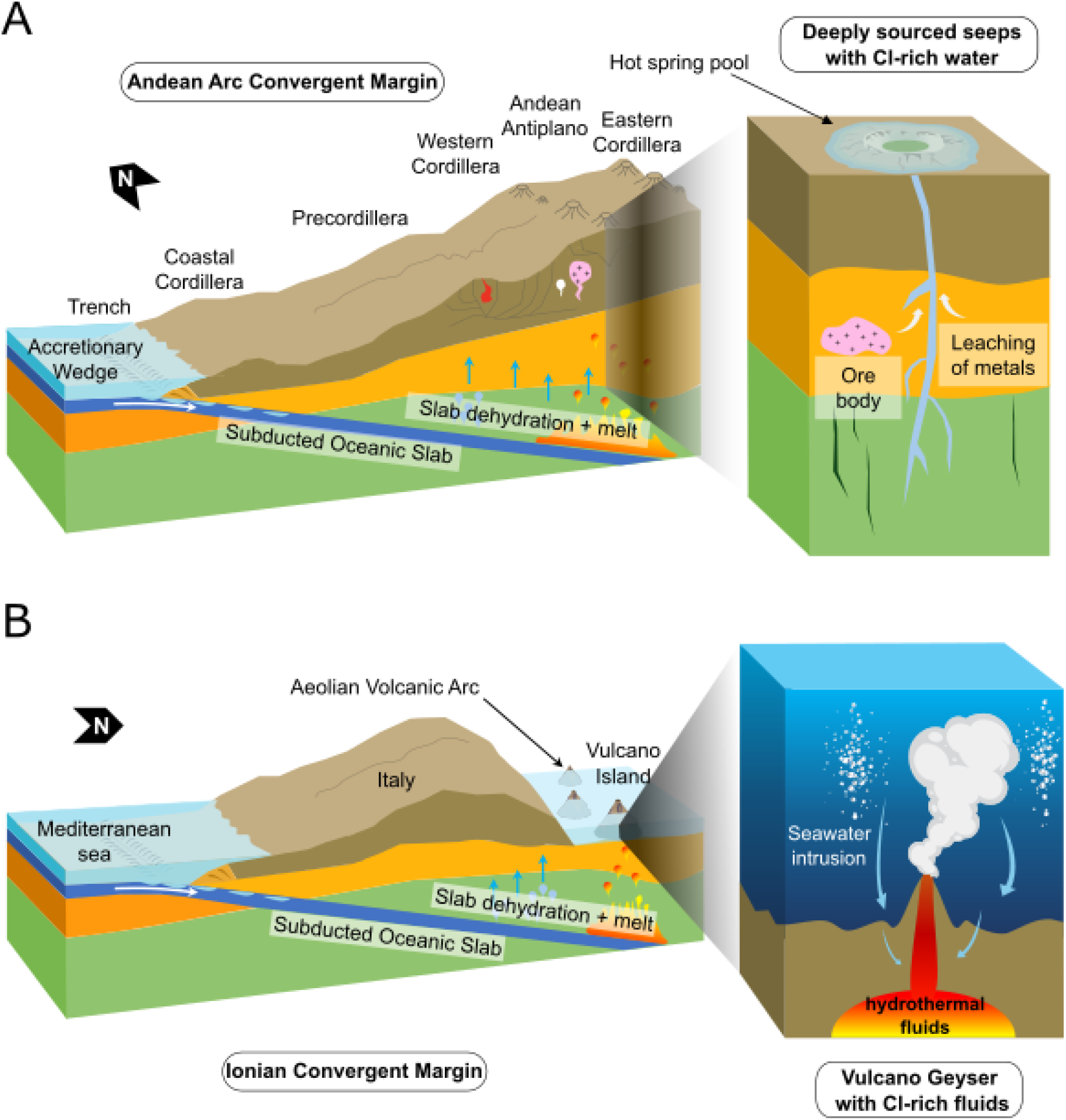
Schematic representation of volcanic arcs at convergent margins and their influence on predicted microbial biosynthetic potential. (a) Schematic representation of the SA-CVZ (South American Central Volcanic Zone) volcanic arc and its influence on microbial biosynthetic potential. On the macro scale, the steeply dipping oceanic slab dehydrates, releasing Cl-enriched fluids from altered oceanic crust. These fluids migrate upward through faults and fractures in the continental crust. Metal leaching in these environments favor NRPS-rich (nonribosomal peptide synthetases) microbial communities. Macro-scale cross-section of the SA-CVZ adapted from ^32^. (b) Schematic of the Ionian convergent margin tectonic system and its role in shaping microbial biosynthetic potential in the AAVP (Aeolian Arc Volcanic Province). The subduction of the Ionian slab beneath the Aeolian Arc fuels arc magmatism and geothermal activity across southern Italy, and enhances mantle input in hydrothermal fluids. Cross-section of the Vulcano Geyser adapted from ^105^.

**Figure 7.**
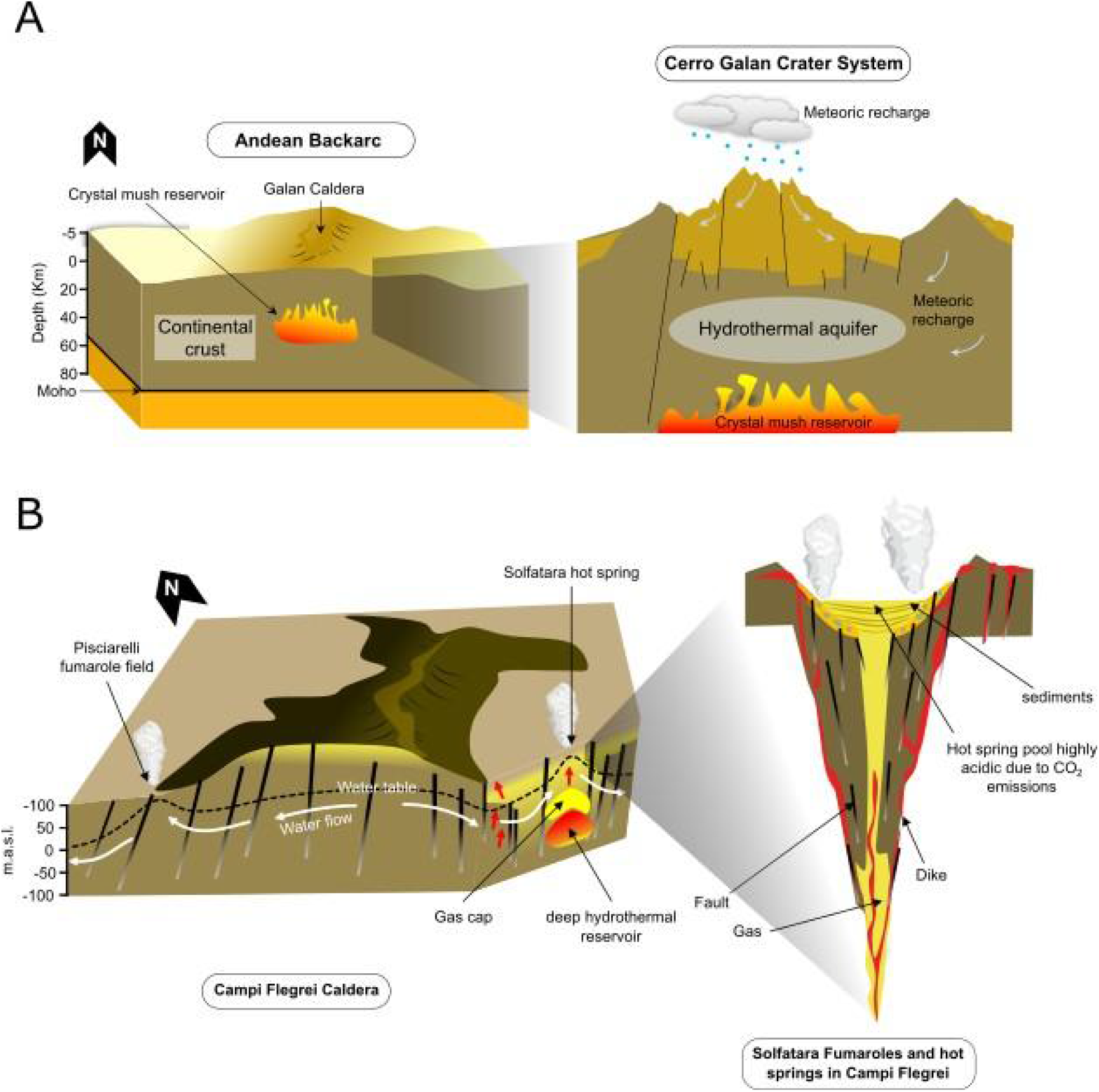
Schematic representation of backarc and post-subduction extensional volcanic systems in active plate tectonic settings and their influence on the predicted microbial biosynthetic potential. (a) Schematic representation of the Galán system in the SA-CVZ (South American Central Volcanic Zone) backarc. Backarc systems are majorly controlled by the hydrothermal circulation of deep hydrothermal reservoirs, receiving less direct input from the subducting slab. Geological and regional scale cartoons were adapted from ^106^. (b) Schematic representation of Campi Flegrei Caldera (post-subduction extensional arc), in the Campania region, Italy; “m.a.s.l.” refers to meters above sea level. Cross sections adapted from ^107^.

In comparison to volcanic arcs, geothermal sites in backarc and post-subduction extensional volcanics show a contraction in putative GCF diversity. This could likely reflect differences in geological context (Figure 7) that shape metal availability. In the SA-CVZ backarc, slab-derived inputs are less pronounced^37^, and the degree of mantle melting and subduction influence is lower than the adjacent volcanic arc^38^. Backarc fluids also interact more extensively with the overriding crust during their ascent, leading to longer residence times^39^. Unlike arcs, they lack abundant direct conduits for magmatic gases to reach the surface, resulting in diminished fluxes of the metal-rich fluids typical of direct slab inputs^39^. By contrast, volcanic arc conduits provide more rapid and direct routes for fluid ascent, delivering a broader mixture of magmatic and hydrothermal fluids^39,40^. The greater fluid diversity in the arc might correspond with the broader predicted GCF diversity we observe, whereas the putative metagenomic GCF profile of backarc sites is accordingly less diverse (Supplementary Figure 2), and specialized in terpenes compared to other active plate tectonic sites (Figure 3).

Backarc geothermal systems are characterized by CO₂-dominated gas compositions (Supplementary Table 3). This environmental setting coincides with the elevated representation of predicted terpene-associated GCFs observed in these systems (Figure 3). While the biosynthesis of some terpene compounds relies on metal cofactors such as Zn, for isopentenyl diphosphate isomerase^41^, and Fe–S clusters for methylerythritol phosphate^42^, their high occurrence in backarc systems could suggest that these pathways might be preferentially maintained for functions beyond metal metabolism. In particular, terpenes can serve versatile roles in membrane fortification^14,43^ and protection against fluctuating pH^44,45^, helping microbial communities withstand CO_2_-rich conditions^46–48^. A comparable trend is observed in the system of the Campi Flegrei, in the Campania region in Italy, where shallow CO_2_ degassing can reach up to 4000–5000 tones per day^49^, and the Solfatara crater and surrounding fumaroles (*e.g.*, Pisciarelli) continuously emit CO_2_-rich, sulfurous fumes^49,50^. Despite such geochemical harshness, cultivation efforts have yielded spore-forming isolates (*e.g.*, *Alicyclobacillus mali* from Pisciarelli mud) whose genomes encode enzymes for unusual isoprenoid terpene biosynthesis^51^. It is worth highlighting this cultivation evidence aligned with our metagenomic data from these Campania systems, which similarly reveals high predicted terpene GCF occurrence (Figure 4). Both the SA-CVZ backarc and Italian Campania cases illustrate that when volcanic systems skew toward high volatile flux and away from direct input of metal-rich magmatic fluids, microbial secondary metabolism might shift toward a greater representation of terpene-associated putative GCFs.

### Potential microbial strategies for metal homeostasis and competition in Mid-Ocean Ridge Ecosystems

Tectonic settings at spreading centers, such as at mid-ocean ridges (MORs), exhibit biosynthetic profiles distinct from both volcanic arcs and backarc systems. In the RVB, geothermal fields like Gunnuhver lie directly atop the Mid-Atlantic Ridge, tapping basaltic magmas and mantle-derived volatiles. At Gunnuhver, subsurface temperatures reach ∼290 °C, with localized zones exceeding 300 °C^52^. Previous studies of Icelandic solfataric fields have reported enrichment in RiPPs, such as archaeocins^53^. Consistent with these observations, RVB hot spring microbiomes in our dataset show the highest representation of predicted RiPP- and NRPS-associated pathways (Supplementary Figure 8), together accounting for 64 % of detected GCF occurrences within this group. The widespread occurrence of putative RiPP-associted GCFs suggests that antimicrobial production might play an important role in these communities: many thermophiles produce heat-stable bacteriocins and related peptides that help maintain competitiveness within dense biofilm assemblages coating mineral deposits in hot springs^12,13^. The measured trace metal compositions of the RVB geothermal include elevated concentrations of Fe, Cu and Zn (Supplementary Table 4), consistent with basalt-hosted hydrothermal fluids generated through fluid-rock interaction in MOR settings^54,55^. However, the bioavailability of these metals can be highly variable, largely because of rapid mineral precipitation, which removes metals from the fluid phase as the fluids ascend and cool^55,56^. In this context, NRPS-associated metallophore pathways may serve a dual role: scavenging essential trace nutrients under limitation^57,58^, and detoxifying excess metals when concentrations become inhibitory^59,60^. The observed putative GCF profile thus can represent a strategy shaped by MOR geochemistry, which creates a complex system for metal bioavailability^54,55,61^.

Overall, our findings suggest that tectonic setting is associated with distinct patterns in microbial GCF composition across geothermal environments. Volcanic arc systems show higher representation of NRPS-associated GCFs, consistent with geochemical conditions influenced by magmatic metal inputs, while backarc systems, especially those located at high elevation, are characterized by comparatively lower GCF diversity, and increased representation of terpene-associated families. In divergent settings such as MOR, the tilt toward NRPS- and RiPP-associated profiles likely reflects strategies for metal scavenging and microbial competition. By influencing magmatic differentiation^62^, pressure of mineral stabilization^63,64^, and fluid-rock interaction pathways^65^, the local tectonic context indirectly influences the chemical and thermal conditions that subsurface microbial communities experience in geothermally influenced ecosystems.

### Microbial terpenes as potential biosignatures in modern and ancient geothermal systems

Beyond differences in putative GCF composition among tectonic settings, the predicted diversity across the sampled sites can provide insights into how geothermal microorganisms have adapted to contrasting geological environments. Among biosynthetic pathways, terpenes represent one of the most chemically diverse classes of microbial natural products^66–68^, making them particularly informative for exploring biosynthetic diversity across geothermal ecosystems. Placement of metagenome-derived sequences alongside reference terpene synthases provides an overview of this predicted diversity (Figure 4). Beyond their ecological significance, microbial terpenes have also attracted interest as many of which hold potential as molecular indicators of past life and environmental conditions^69^. Although most biomarker applications to-date rely on plant-derived terpenes (e.g., triterpenoids) preserved in sedimentary archives from the mid-Paleozoic onwards^70,71^, recent work has highlighted the presence of microbial terpenes in modern and ancient hydrothermal deposits^69,72^.

Notably, SKVP geothermal sites contained the highest proportion of putative terpene-associated GCFs, almost double that observed in tectonically active convergent margins at volcanic arcs, (Figure 3c, Supplementary Figure 9). High terpene occurrence and volatile organic compounds have also been documented in the Outokumpu deep drill site in Finland^73^, which is hosted within a paleoserpentinized ultramafic complex, emplaced and metamorphosed during the Svecofennian orogeny^74^. The recurrence of terpene-dominated signatures in inactive plate tectonic locations either plume-related (*e.g* SKVP), or ancient intraplate settings (*e.g* Outokumpu) might suggest that long-lived hydrothermal circulation with limited magmatic metal inputs can foster modern microbial communities that preferentially invest in membrane-protective and stress-mitigating terpene pathways. Such parallels point to the potential persistence of terpene biosynthetic strategies in hydrothermal systems over geological timescales, as terpene signatures could be constantly present even if plate tectonics ceases.

The elevated representation of predicted terpene-associated GCFs in the SKVP plume-related geothermal systems contrasts with the active convergent margin tectonics of volcanic arcs, where putative NRPS-associated GCFs are more prevalent (Figure 3c). In arc settings such as the SA-CVZ, CAVA, and AAVP, hydrothermal fluids are directly influenced by magmatic degassing^75^, and subducting slab metal input^76^. Under these conditions, NRPS systems provide a selective advantage by producing metal-binding metabolites that mitigate toxicity^77,78^. Similar high occurrence of NRPS-associated families is observed in other tectonically active volcanic settings, such as the Kolumbo submarine volcano (Santorini)^79^, shallow-water hydrothermal vents in the Azores^80^, and the Barren Islands volcanic arc in Southeast Asia^81^. In contrast, the SKVP hot springs are primarily fed by meteoric waters circulating through the continental crust, with limited magmatic input but constant heat-flux from remnants of the intraplate plume (Figure 8). Here, other environmental factors, such as the local thermal regime and the arid, high altitude desert like nature of the region, might favor the protective and stabilizing functions of terpenes^14,82^. These compounds are well-suited to mitigate oxidative stress^83,84^, maintain membrane integrity under variable temperature and pH conditions^85–87^, and enhance UV protection^88^. Thus, intraplate tectonic systems hosting residual geothermal activity may have the potential to foster microbial communities that harbor terpene-focused strategies of stress resilience, in contrast to the NRPS-associated metal response typical of volcanic arcs in active convergent margins of subduction zones (Figure 9).

**Figure 8.**
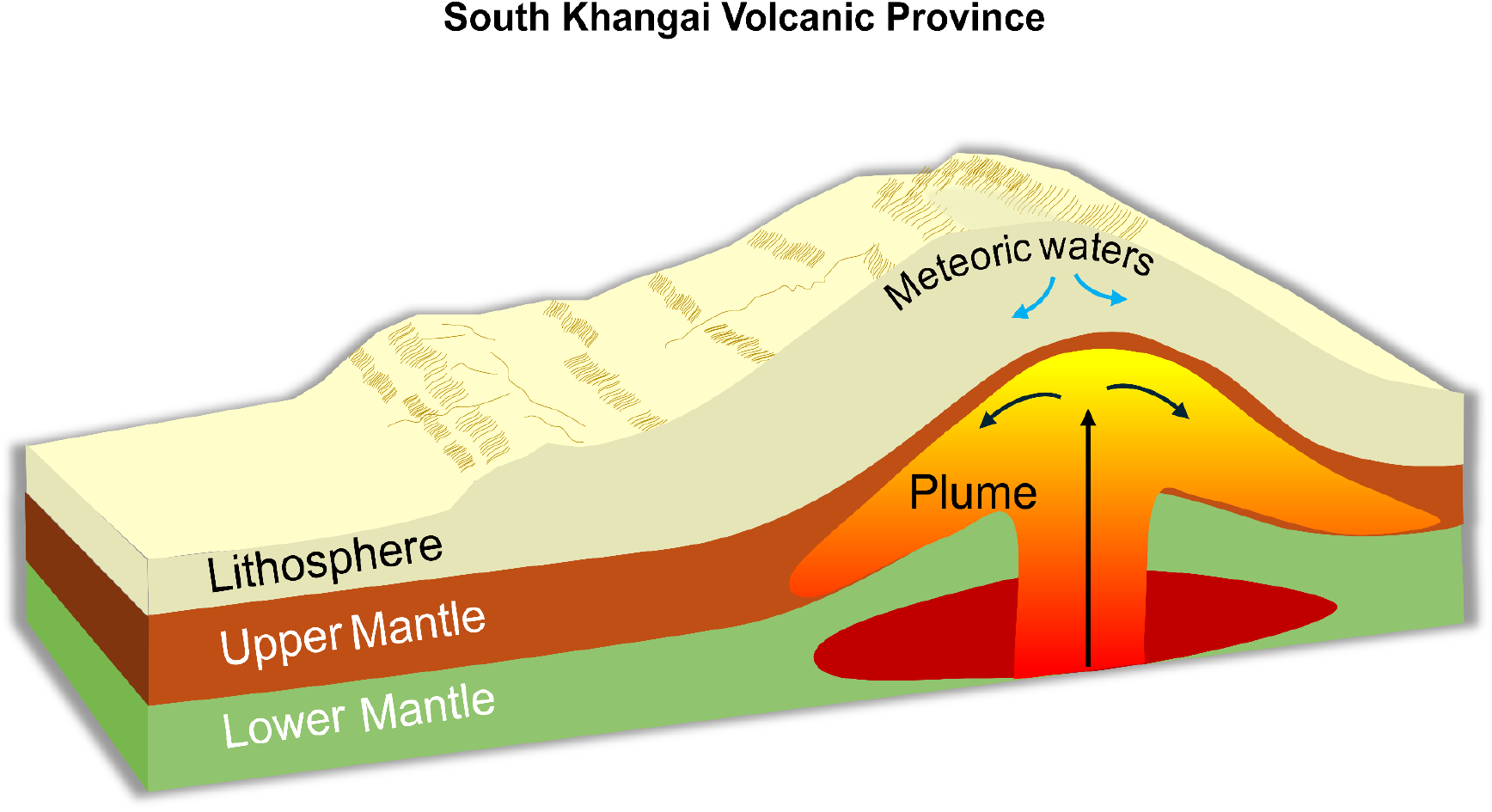
Geological context and distribution of biosynthetic gene cluster families (GCFs) across the South Khangai Volcanic Province (SKVP). (a) Conceptual cross-section of the plume–lithosphere interaction in the SKVP, modified from ^108^.

**Figure 9.**
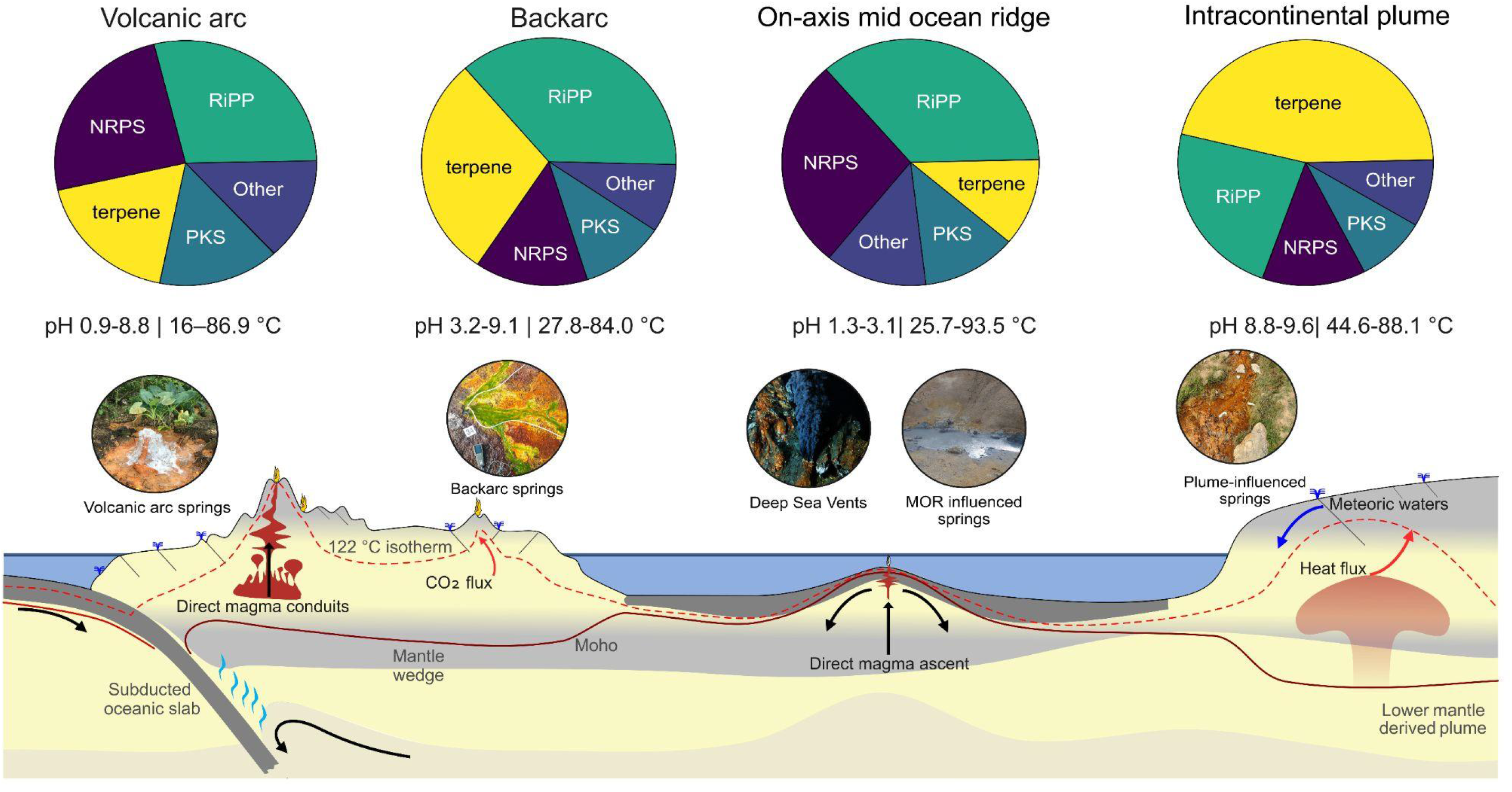
Schematic cross-section of representative tectonic regimes and associated geothermal systems in this study, illustrating large-scale geodynamics shaping microbial biosynthetic gene cluster family diversity. In volcanic arcs, fluids sourced from subducting slabs and magmatic degassing might support NRPS-linked (nonribosomal peptide synthetases). Backarc springs, with reduced direct magmatic input, could shift toward stress-adaptive terpene pathways. Mid-ocean ridge (MOR) systems, might display NRPS- and RiPP-linked (ribosomally synthesized and post-translationally modified peptides) repertoires. Intraplate plume-influenced settings might be characterized by high presence of terpene GCFs potentially reflecting scarce direct magmatic input in this system. GCF charts represent the relative proportions of GCF categories within the SA-CVZ (South American Central Volcanic Zone) volcanic arc, SA-CVZ backarc, RVB (Reykjanes Volcanic Belt) and SKVP (South Khangai Volcanic Province) respectively. Moho and 122°C isotherm are representative general changes across tectonic settings. Cross section adapted from ^89^.

## Conclusion

Understanding how geological and geochemical gradients relate to potential microbial secondary metabolism across large planetary scales provides insight into the ecological organization of microbial communities in extreme environments. Here, we show that large-scale geological context is associated with distinct patterns in biosynthetic GCF composition across globally distributed geothermal systems. Volcanic arc microbiomes are characterized by a greater presence of putative NRPS-associated GCFs, whereas backarc, post-subduction extensional, and intraplate plume-related geothermal systems show increased representation of putative terpene-associated GCFs. We further demonstrate that geothermal microbiomes harbour extensive biosynthetic novelty, with most predicted GCFs lacking similarity to experimentally characterized biosynthetic pathways, highlighting these environments as a substantial reservoir of unexplored natural product diversity. Together, these results suggest that tectonic and geochemical settings provide an important, yet previously underexplored, framework for interpreting variation in microbial biosynthetic signatures and provide geological context for future natural product discovery and might provide a new framework for reconstructing ancient environments on early Earth and beyond.

## Methods

### Sample acquisition and locations

The present work encompasses sampling campaigns spanning different tectonics settings distributed worldwide (Supplementary Methods, Supplementary Table 6), and following a large-scale sampling approach described in ^89^ (Figure 1), that allows the use of deeply-sourced seeps to access information into the subsurface biosphere^115^. In this work, samples from deeply-sourced seeps and other geothermal features were used together with geothermal biofilm and sediments as representative of geothermally influenced sites. Environmental metadata and geological context were integrated based on the extended geochemical and tectonic framework outlined in ^89^, which emphasizes the role of deep subsurface inputs in structuring microbial communities. This broader context allowed us to systematically categorize geothermal sites by their underlying tectonic regime (backarc, volcanic arc, on axis ridges, post-subduction extensional arcs and intraplate plume systems), facilitating standardized comparisons across disparate geographic regions.

Briefly, at each site, the primary thermal feature (i.e., venting point) was identified based on temperature, pH and conductivity gradients in combination with visual cues (e.g., gas bubbling, mineral precipitates), as described by ^89^. Microbial cells were retrieved from fluids collected directly from the venting orifice using a titanium pipe to avoid air and metal contamination, sampled through a silicone tubing attached to a portable peristaltic pump, and filtered through Sterivex 0.22 μm cartridges (MilliporeSigma). At least 2 L of hydrothermal fluid was filtered at each site and filters stored at –20 °C on-site, except for the SA-CVZ volcanic arc and backarc where samples were transported in liquid nitrogen dry shippers. Sediment samples were collected in close proximity (within centimeters) to the fluid vents using sterile, acid-washed PTFE spatulas and transferred into 50 mL centrifuge (i.e., Falcon) tubes. Sediment aliquots were also sampled for subsequent analysis of trace elements concentration and CNS content in each site. Approximately 40–45 mL of sediment was transferred into 50 mL centrifuge tubes. Collected sediment samples were immediately frozen at –20 °C and kept at this temperature during transport to the laboratory, where they were stored until further analyses. Gas-phase and water samples for dissolved gases were collected in pre-evacuated 250 mL Giggenbach bottles containing 50 mL of 4 N NaOH, following the method of ^90^, to minimize atmospheric contamination. Additionally, copper tubes were used to sample noble gases at each site, as detailed by ^90^.

Physicochemical parameters of the fluids collected to provide environmental context to the sampled sites were measured *in situ* using a thermocouple and a multiparameter probe (HANNA, HI98196). The thermocouple was used to record pool temperatures directly at the main vent inlet. Freshly venting fluids were analyzed with the multiprobe, which measured pH, oxidation-reduction potential, specific conductivity, dissolved oxygen, and total suspended solids. For fluid temperatures exceeding 55 °C, corresponding to the maximum operating limit of the dissolved oxygen sensor, samples were cooled in closed containers prior to measurement to allow analysis within instrument specifications. Cooling was performed under conditions minimizing gas exchange with the atmosphere. The impact of this procedure was evaluated through laboratory tests of dissolved oxygen measurements obtained before and after controlled cooling, which indicated that deviations remained within the analytical uncertainty of the sensor under our measurement conditions.

The dataset consisting of 216 metagenomic assemblies analyzed in this study was supplemented using 3 additional assemblies publicly available on NCBI and representing the Guaymas basin, totalling 219 samples. These samples were selected based on (i) comparable environmental context (hydrothermal/geothermal sediments), and (ii) availability of raw sequencing data. Collection strategy for deep-sea hydrothermal vent microbial biomass at the East Pacific Rise (EPR; 9°50 N, 104°17 W) was described in ^91^. Briefly, a total of nine samples were recovered using both experimental microbial colonization devices deployed by the ROV Jason and the deep-submergence vehicle Alvin. Samples were collected from Crab Spa (9°50.39’ N, 104°17.48’ W: 2503-m depth), P-vent (9°50.28’ N, 104°17.47’ W: 2506-m depth), Teddy Bear (9°50.50’ N, 104°17.51’ W: 2514-m depth), Bio9 (9°50.30’ N, 104°17.30’ W: 2503-m depth), Tica (9°50.39’ N, 104°17.49’ W: 2511-m depth), Perseverance (9°50.95’ N, 104°17.59’ W: 2507-m depth).

### DNA extraction, metagenomic sequencing and BGC analysis

DNA was extracted with a combination of the DNeasy PowerSoil Kit (QIAGEN), and a modified phenol-chloroform DNA extraction method adapted for low biomass samples ^19^, to maximize DNA recovery across heterogeneous sample types and low-biomass environments. Sequencing was carried out by Novogene (UK) using the NGS DNA Library Prep Set (Cat No.PT004) and Illumina platform NovaSeq X Plus. Briefly, genomic DNA was randomly sheared into short fragments, which were then end repaired, A-tailed, and ligated to Illumina adapters. Adapter-ligated fragments were size selected, purified, and processed according to Novogene’s standard workflow prior to sequencing. Libraries were quantified using Qubit and qPCR, and fragment size distributions were assessed prior to pooling and sequencing. Sequencing quality control and read filtering were performed by Novogene, including removal of adapter-contaminated reads, reads containing >10% ambiguous bases (N), and reads containing >50% low-quality bases (Q ≤ 5). Per-sample sequencing depth and statistics are reported in Supplementary Table 7.

The GEOMOSAIC pipeline (github.com/giovannellilab/Geomosaic, Corso et al. 2025, *submitted*), designed to integrate state of art metagenomic software analysis with environmental data, was used for the sequences to ensure uniformity of the data. Briefly, all the raw reads were trimmed using fastp, set to default parameters. The trimmed reads were then assembled using metaSPAdes (v. 3.15.5)^92^ with k-mers -k 33, 55, 77, and 127, and filtering for minimum contig length of 2000 base pairs. All the generated 219 assemblies were analyzed with antiSMASH (antibiotics and Secondary Metabolite Analysis SHell) 7.0^93^, using default settings, including KnownClusterBlast/MIBiG (Minimum Information about a Biosynthetic Gene cluster) 4.0^94^ matching and the product/substrate prediction modules to investigate the richness of biosynthetic potential in geothermal environments. AntiSMASH detects BGC classes by rule-based domain profiling and, when possible, predicts candidate product scaffolds. To account for redundancy among similar BGCs predicted across multiple assemblies, antiSMASH GenBank outputs were dereplicated into gene cluster families (GCFs) using BiG-SCAPE (Biosynthetic Gene Similarity Clustering and Prospecting Engine) 2.0^95^. BiG-SCAPE computes pairwise similarity relationships among BGCs based on domain architecture, sequence similarity, and gene organization, generating similarity networks in which nodes represent individual BGCs and edges represent similarity scores.

GCFs were defined using a similarity cutoff of 0.5 following standard BiG-SCAPE clustering parameters. Analyses were performed separately for major biosynthetic classes (NRPS, PKS, RiPPs, and terpenes) and included reference BGCs from the MIBiG (Minimum Information about a Biosynthetic Gene cluster) database^126^. Within each similarity network, connected components were interpreted as gene cluster families, thereby grouping homologous or partially fragmented BGCs originating from different assemblies into unified biosynthetic units.

### BGC dereplication and novelty assessment

Biosynthetic profiles were represented as presence–absence matrices of GCFs per metagenome. No abundance estimates were inferred and read mapping was not performed. Analyses therefore focus on biosynthetic diversity rather than quantitative abundance, to ensure comparability across samples with heterogeneous sequencing depth and assembly completeness.

A GCF was considered present in a sample when at least one member BGC belonging to that family was detected within the corresponding assembly. Because metagenomic assemblies may contain incomplete or fragmented BGCs, dereplication at the GCF level reduces redundancy by clustering partial and complete clusters sharing conserved biosynthetic architecture. Analyses were therefore conducted at the family level rather than individual BGC predictions. GCFs detected in fewer than 5% of samples were removed to reduce sparsity. Sequencing depth (log₁p-transformed read counts) was included as a conditioning variable in multivariate analyses to control for technical variation. To assess whether sequencing depth influenced biosynthetic profiling and clustering, the number of reads per metagenome (Supplementary Table 7) was used as a proxy for sequencing effort. Initial ordination analyses showed significant correlations between sequencing depth and ordination axes (Spearman ρ = −0.68, p < 2.2 × 10⁻^16^ for Axis 1; ρ = −0.28, p = 2.3 × 10⁻^5^ for Axis 2), indicating a potential technical gradient. To control for this effect, sequencing depth was log-transformed (log1p) and included as a conditioning variable in constrained ordination models (capscale). After conditioning on sequencing depth, correlations between ordination axes and sequencing depth were no longer significant (Axis 1: ρ = −0.04, p = 0.51; Axis 2: ρ = 0.005, p = 0.94).

Results were combined into a single feature table and imported in R (v. 4.3, R Core Team 2024). The subsequent statistical analyses, data processing and plotting were carried out using the phyloseq ^96^, vegan^97^ and ggplot2^98^ packages. Alpha diversity of GCFs was quantified using Shannon’s diversity index, calculated from BiG-SCAPE-derived GCF profiles. Differences among tectonic settings (volcanic arc, backarc, post-subduction) were tested with the Kruskal–Wallis test followed by pairwise Wilcoxon rank-sum tests with Benjamini–Hochberg correction. Relationships between environmental variables (e.g., temperature, pH, conductivity, crustal thickness) and biosynthetic profiles were assessed using partial redundancy analysis (RDA) conditioned on sequencing depth, with significance evaluated using permutation tests (999 permutations). Prior to analysis, conductivity (spc), chloride, and sulfate were log₁₀-transformed to reduce skew. Sample clusters were identified using k-means clustering of RDA site scores (k = 3), selected based on elbow and silhouette criteria. While silhouette width was maximized at k = 2, the elbow criterion indicated diminishing returns beyond k = 3, and k = 3 provided improved ecological interpretability and separation of major geothermal system types (Supplementary Figure 10). Differences among clusters were evaluated using PERMANOVA and ANOSIM, and homogeneity of dispersion was verified using betadisper (p = 0.31), indicating that separation reflected compositional differences rather than unequal variance among clusters. Finally, differences among clusters in individual environmental variables were tested using the non-parametric Kruskal–Wallis test with Dunn’s *post hoc* comparisons and Benjamini–Hochberg correction.

### Phylogenetic analysis of core terpene biosynthetic proteins

To characterize putative terpene biosynthetic diversity, we performed a phylogenetic analysis of predicted core terpene biosynthetic proteins recovered from antiSMASH-annotated biosynthetic gene clusters. To reduce redundancy, a single representative environmental BGC was selected from each terpene-associated gene cluster family (GCF) identified by BiG-SCAPE prior to phylogenetic analysis, while reference terpene biosynthetic proteins from MIBiG were retained to provide phylogenetic context. AntiSMASH GenBank output files (.gbk) generated from metagenomic assemblies were parsed using a custom Python script implemented with Biopython^99^ (v 1.71). Coding sequences (CDS features) were screened for terpene-associated annotations contained within the gene_functions and sec_met_domain qualifier fields produced by antiSMASH. CDS entries were retained as candidate terpene core biosynthetic genes when annotation text contained terpene-related functional keywords corresponding to terpene synthases or associated prenyltransferases (including: *phytoene_synt*, *Lycopene_cycl*, *terpene_cyclase*, *Terpene_synth*, *Terpene_synth_C*, *trichodiene_synth*, *NapT7*, *TRI5*, *fung_ggpps*, and *fung_ggpps2*).

For each representative environmental BGC, the translated amino acid sequence highest-ranking candidate terpene core biosynthetic protein was extracted. Candidate core proteins were prioritized according to antiSMASH functional annotations, protein completeness, and BGC completeness to preferentially retain complete terpene synthases or cyclases when multiple candidates were present within the same BGC. FASTA headers encoded sequence provenance, including source GenBank file, record identifier, locus tag, and antiSMASH functional annotations. Extracted protein sequences from metagenomic assemblies were combined with reference terpene biosynthetic proteins derived from curated biosynthetic gene clusters to provide phylogenetic context.

Protein sequences were aligned using MAFFT^100^ (v7; with the flags “--auto”). Alignment trimming was performed using ClipKIT^101^ (v 2.3.0) with default parameters to remove poorly aligned positions and reduce noise while retaining phylogenetically informative sites. Maximum-likelihood phylogenetic trees were inferred using IQ-TREE 2^102^ under the LG+G4+F amino acid substitution model. Branch support was evaluated using 1,000 ultrafast bootstrap replicates and 1,000 SH-like approximate likelihood ratio tests (SH-aLRT). Phylogenetic tree display and annotation was performed through iTOL^103^ v7. The whole pipeline was run on the HPC IBiSco (Infrastructure for BIg data and Scientific COmputing; project PON R&I 2014-2020 dell’ Avviso 424/2018 - Azione II. 1.) at the University of Naples Federico II, using SLURM^104^ (v 2.11.5).

## Conflicts of Interest Statement

The authors declare that the research was conducted in the absence of any commercial or financial relationships that could be construed as a potential conflict of interest.

## Data and code availability

Raw sequences are available through the European Nucleotide Archive (ENA) under the Umbrella Project CoEvolve PRJEB55081. The NCBI samples from EPR are available through the NCBI PRJ PRJNA1092253. The NCBI samples from the Guaymas basin are available through the NCBI PRJ number PRJNA879229. A complete shell script and jupyter notebook containing all the steps to reproduce our antiSMASH pipeline and figures in this paper is available at: github.com/giovannellilab/antismash_BGC_geothermal_env with DOI https://doi.org/10.5281/zenodo.15785034 along with all collected geochemical and environmental data.

## Acknowledgements

The authors wish to thank all the personnel that participated in the field campaigns and helped to collect the samples which lead to the metagenomic libraries used in this work. We express our gratitude to the “Secretaría de Medio Ambiente de la Provincia de Catamarca” for their administrative support that allowed our sampling (Expte.EX-2022-02222431-CAT-DPB#SEAS, Resolución S.E.A., D.S. N°: 053/2017). Additionally, we thank the “Secretaría de Política Ambiental en Recursos Naturales, Ministerio de Ambiente y Desarrollo Sostenible de la Nación Argentina” for providing the certificate of compliance (Title: IF-2023-65966642-APN-SPARN#MAD, UId: ABSCH-IRCC-AR-264943-1) and the export certificate for genetic resources (CE-2023-69835755-APN-SPARN#MAD). We also appreciate the support of Corporación Nacional Forestal (CONAF, Chile). We thank the native communities of Antofalla, El Peñón, Antofagasta de la Sierra and Colla communities for their support during fieldwork and access to sites. Additionally, we thank the Environment and Energy Agency of Iceland (license n. UST202305-096/D.Ö.H. and OS2023010131/50.4.3), the Department of Environment and Tourism of Uvurkhangai aimag of Mongolia (permit n. 144 of 2023.07.26).

## Author Contributions

Conceptualization: A.C.P.S., B.dP., D.G. Methodology: All authors. Data curation: A.C.P.S., B.dP., F.M., M.C., D.B., B.B., M.S., C.V., A.V.B., P.B., D.G. Visualization: A.C.P.S., B.dP., B.B., D.G. Investigation: All authors. Code: A.C.P.S., B.dP. Funding: D.G.. Writing - original draft: A.C.P.S., B.dP., D.G., Writing - review & editing: All authors.

## CoEvolve Project Consortium

Donato Giovannelli, CoEvolve consortium PI, Department of Biology, University of Naples “Federico II”, Naples, Italy

Angelina Cordone, Department of Biology, University of Naples “Federico II”, Naples, Italy

Agostina Chiodi, CONICET Instituto de Bio y Geociencias del NOA (IBIGEO), Rosario de Lerma, Argentina

Alberto Vitale Brovarone, Dipartimento di Scienze Biologiche, Geologiche e Ambientali - BiGeA, Università degli Studi di Bologna Alma Mater, Bologna, Italy

Alessia Bastianoni, Department of Biology, University of Naples “Federico II”, Naples, Italy Annarita Ricciardelli, Department of Biology, University of Naples “Federico II”, Naples, Italy Bernardo Barosa, Department of Biology, University of Naples “Federico II”, Naples, Italy Carlos Ramirez Umana, Servicio Geológico Ambiental (SeGeoAm), Heredia, Costa Rica Davide Corso, Department of Biology, University of Naples “Federico II”, Naples, Italy Deborah Bastoni, Department of Biology, University of Naples “Federico II”, Naples, Italy Edoardo Taccaliti, Department of Biology, University of Naples “Federico II”, Naples, Italy

Federico A. Vignale, European Molecular Biology Laboratory - Hamburg Unit, Hamburg, Germany

Feliciana Oliva, Department of Biology, University of Naples “Federico II”, Naples, Italy Flavia Migliaccio, Department of Biology, University of Naples “Federico II”, Naples, Italy

Francesco Montemagno, Department of Biology, University of Naples “Federico II”, Naples, Italy

Gabriella Gallo, Department of Biology, University of Naples “Federico II”, Naples, Italy

Gerdhard L. Jessen, Instituto de Ciencias Marinas y Limnológicas, Universidad Austral de Chile, Valdivia, Chile

J. Maarten de Moor, Observatorio Vulcanológico y Sismológico de Costa Rica (OVSICORI), Universidad Nacional, Heredia, Costa Rica

Jacopo Brusca, Department of Biology, University of Naples “Federico II”, Naples, Italy

James A. Bradley, Aix Marseille Univ, Université de Toulon, CNRS, IRD, MIO, Marseille, France

Karen G. Lloyd, Earth Science Department, University of Southern California, Los Angeles, CA, USA

Luca Tonietti, Department of Science and Technology, Parthenope University of Naples, Naples, Italy

Luciano di Iorio, Department of Biology, University of Naples “Federico II”, Naples, Italy

Maria García Alai, European Molecular Biology Laboratory - Hamburg Unit, Hamburg, Germany

Martina Cascone, Department of Biology, University of Naples “Federico II”, Naples, Italy

Matteo Selci, Department of Biology, University of Naples “Federico II”, Naples, Italy

Mattia Esposito, Department of Biology, University of Naples “Federico II”, Naples, Italy

Monica Correggia, Department of Biology, University of Naples “Federico II”, Naples, Italy

Mustafa Yucel, Institute of Marine Sciences, Middle East Technical University, Erdemli, Turkey

Nunzia Nappi, Department of Biology, University of Naples “Federico II”, Naples, Italy

Peter H. Barry, Marine Chemistry & Geochemistry Department, Woods Hole Oceanographic Institution, MA, USA

Sara Claudia Diana, Department of Biology, University of Naples “Federico II”, Naples, Italy

Stefano Caliro, Osservatorio Vesuviano, Istituto Nazionale di Geofisica e Vulcanologia (INGV), Napoli, Italy

## Funding

This work was supported by funding from the European Research Council (ERC) under the European Union’s Horizon 2020 research and innovation program Grant Agreement No. 948972—COEVOLVE—ERC-2020-STG. ACPS was supported by the CRESCENDO DP Marie Skłodowska-Curie programme (MSCA-COFUND-2020) with Grant Agreement No. 101034245. BdP was supported by the Marie Skłodowska-Curie Actions Postdoctoral Fellowship (Grant Agreement No. 101154017; HORIZON-MSCA-2023-PF-01 project SUBCARB) under the European Union’s Horizon Europe programme. FM was supported by the “National Biodiversity Future Center-NBFC,” project code CN_00000033, concession decree no. 1034 of 17 June 2022 adopted by the Italian Ministry of University and Research. Additional support came from The National Fund for Scientific and Technological Development of Chile (FONDECYT) Grant 1251543 (The National Research and Development Agency of Chile, ANID Chile), and COPAS COASTAL ANID FB210021 to G.L.J. PHB acknowledges NSF award 2121637, which partially supported this work. CV acknowledges NSF awards DEB 1136451 and OCE 1948623.

## Notes

https://github.com/giovannellilab/antismash_BGC_geothermal_env

https://doi.org/10.5281/zenodo.15785035

https://www.ebi.ac.uk/ena/browser/view/PRJEB55081

https://www.ncbi.nlm.nih.gov/bioproject/PRJNA879229/

### Competing Interest Statement

The authors have declared no competing interest.

### Summary of Updates

Figures 3-9 revised, metadata table added

