## Supplementary Materials for "Tectonic setting shapes microbial biosynthetic potential across global geothermal environments"

### Sample locations

#### Reykjanes Volcanic Belt (RVB)

We collected 10 samples from 3 geothermal spring regions in Krýsuvík (63.895451; -22.056914), Gunnuhver (63.819344; -22.68), and Hveragerdi (64.008117; -21.17949). The sampling areas were characterized by absence of vegetation, and hot soil located both to the north and south of the main vent area. The soil type was predominantly clayey/clayey-sandy.

#### South Khangai Volcanic Province (SKVP)

We collected a total of 7 samples from 4 geothermal springs in the regions of Övörkhangai and Bayankhongor. The Khujirt seep (46.9019233; 102.7696788) was surrounded by anthropic structures and tubings, probably placed to collect the water. The Shargaljuut spring (46.33309; 101.224945) was located near Erdenetsogt village, within the Bayankhongor Province. This was a popular area for locals, known for the hot springs with beneficial properties. The Nariinteel (46.128628; 101.605386) and Taragt (46.273908; 102.473959) springs were located in Övörkhangai in flat areas with low vegetation.

#### South American Central Volcanic Zone (SA-CVZ)

We collected a total of 68 samples from 50 geothermal spring locations in the SA-CVZ. The full expedition report to the SA-CVZ is present in [1]. Briefly, deeply-sourced fluids were collected in the Andean Convergent Margin, between 17 °S and 24 °S, to understand interactions between microbiology, deeply-sourced fluids, the crust, and tectonic parameters.

#### Central America Volcanic Arc (CAVA)

We collected a total of 88 samples from 56 geothermal spring locations in the CAVA . Full details of sampling are described in [2]. Briefly, samples were located along the southern CAVA, to investigate whether geological setting and context could influence subsurface microbiology.

#### Tuscan-Latium Volcanic Province (TLVP)

We collected a total of 7 samples from 7 geothermal springs in the TLVP. Sorgente di Chiorba spring (43.1527486; 10.853645) emerges from an opening located in a stone made structure located in the Biancane natural park; Sasso Pisano (43.1676069; 10.8667609) is a degassing site located close to a camping area; Soffioni Boraciferi (43.1435272; 10.8153437) is situated within a large bubbling pool in an area with very little vegetation; Torre Alfina (42.747778; 11.948202) is a pool of meteoric water with diffuse degassing; Solfatara di Ferento (42.504311; 12.1371561) is a site of diffuse degassing within a river margin; San Cristoforo (42.3984606; 12.0592911) is a pool of hot water carved in a carbonate mound; Stagno Bianco (41.7045854; 12.5365787) is a spring in a small canyon located in a degassing area.

#### Campania

We collected a total of 17 samples from 12 geothermal springs in the Campania region. Bagnone (43.181228; 10.832435); Acque Cantani (41.314806; 13.891273), Terme Caracciolo Forte (40.690045; 15.251482); Capasso geyser (40.690045; 15.251482), Sorgente Ferrata (40.651557; 15.229379); Grotta dell'acqua (40.825108; 14.059695); Lido lo scoglio (40.82538; 14.077299); Madonna dei Lattani (41.302894; 13.984734); Piccolo Inferno (41.314487; 13.894006); Stufe di Nerone (40.826908; 14.076022); Sorgente Petrinum (41.125096; 13.890296); Varchera (40.705666; 15.222618).

#### Aeolian Arc Volcanic Province (AAVP)

We collected a total of 10 samples from 10 geothermal springs in the AAVP. Acque Calde (38.417502; 14.959262) is a pool characterized by diffused degassing located in the island of Vulcano; Fangaia Vulcano (38.416171; 14.959509) is a mud pool which has been previously geochemically characterized[3]; Levante Bay (38.4167042; 14.9599477), Geyser Vulcano (38.4175871; 14.9599809), Black Point (38.3814; 15.0618), Bottaro (38.3819; 15.0637) are shallow-water hydrothermal vents; Hotel Oasi (38.637719; 15.075066) is located inside the touristic Hotel Oasi in Panarea where hydrothermal waters come from a well; Bagno Secco Sorgente Rossa (38.491391; 14.908525) is a spring located in an area with abundant vegetation; Terme San Calogero Stufa (38.477998; 14.910223) is located in an old thermal stove in the thermal establishment San Calogero in Lipari; Casa Fulco (38.798403; 15.237873) is a water well constantly monitored by the National Volcanology and Geophysics Institute in Italy (INGV).

### Geochemical analysis

The ionic composition of the geothermal fluids were investigated using ion chromatography (IC) on filtered water samples using a Metrohm ECO IC system, following the standardized procedures detailed in [4]. In the field, fluids were filtered through 0.22 μm membrane filters and collected in acid-washed plastic containers. To prevent contamination and to ensure analytical accuracy, both filters and collection vessels were rinsed with sample fluid three times prior to use. Samples were stored at 4 °C and processed within recommended holding times depending on target analytes ($Cl^-$, $Br^-$, $NO_3^-$, $NO_2^-$, $SO_4^{2-}$, $PO_4^{3-}$, $Ca^{2+}$, Na, K, $Mg_2^+$, $NH_4^+$). For chloride preservation, ethylenediamine (EDA) was added in accordance with [5], when applicable. Cation and anion analyses were performed in separate runs using Metrosep C 4 and A Supp 5 columns, respectively. Samples were diluted with ultrapure water (typically 1:10) to maintain conductivity below 600 µS/cm, which was critical for optimal chromatographic resolution and column longevity. Additional pre-treatment steps were applied selectively based on sample composition: silver ion exchange was used in samples with elevated chloride concentrations to reduce interference with anion detection, while solid-phase extraction (C18) was applied to samples with high dissolved organic compounds to improve chromatographic resolution[4].

Chromatographic separation was achieved using a guard column, followed by the analytical column and conductivity detector. For anion analysis, chemical suppression was applied to reduce background conductivity and enhance analyte signals. Compounds were identified based on retention times matched against certified standards, and quantification was performed through multipoint calibration curves (0.1–10 mg $L^{-1}$). Calibration linearity was assessed using the coefficient of determination ($r^2$), and only curves with $r^2 \geq 0.999$ were accepted. Calibration was verified by periodic analysis of a 1 ppm multistandard every 10 samples to monitor instrument stability and peak drift[4].

All samples were analyzed in triplicate.Quality control included routine analysis of blanks to verify low background signal and absence of carryover, as well as standard reference solutions to assess analytical accuracy, with acceptable recoveries in the range of 80–120%[4]. This workflow allowed precise quantification of environmentally relevant ions (*e.g.*, sulfate, chloride, sodium, potassium, calcium, magnesium), facilitating site comparisons and enabling downstream ecological interpretations[4].

Concentrations of trace elements were determined by inductively coupled plasma mass spectrometry (ICP–MS) following the protocols described by [6]. Briefly, filtered and acidified hydrothermal fluid samples were analyzed on an Agilent 7900 ICP–MS equipped with a collision/reaction cell to minimize polyatomic interferences. Calibration was carried out using multi-element standards spanning 0.01–100 µg $L^{-1}$, with internal standards (e.g., Rh, In) added to correct for instrumental drift. Accuracy and precision were verified against certified reference materials and matrix-matched standards, ensuring reliable quantification even in high-salinity geothermal fluids. The procedure allows detection of trace metals of biological relevance (e.g., Fe, Mn, Co, Ni, Cu, Mo, W, V, As) at sub-ppb levels.

Gas composition analysis was performed at the Volcano Observatory of the Universidad Nacional de Costa Rica. Headspace gases (He, $H_2$, $O_2$, Ar, $N_2$, $CH_4$) were analyzed using an Agilent 7890a gas chromatograph equipped with two HP-molesieve columns (Agilent 19095P-MSO) maintained at 30 °C. Methane was detected with a flame ionization detector, while the remaining gases were quantified using thermal conductivity detectors. Following gas chromatograph analysis, the NaOH solution from the Giggenbach bottles was titrated with 0.1 N HCl to determine $CO_2$ content. The $CO_2/CH_4$ ratio was calculated by determining the total moles of $CO_2$ and $CH_4$ in each sample. Carbon isotope compositions ($\delta^{13}C$) of gas samples were measured using a Picarro G2201-I analyzer, following acidification of the NaOH solutions extracted from the Giggenbach bottles. $\delta^{13}C$ values (reported in ‰ relative to the PDB standard) were calibrated against a set of eight reference standards ranging from +2.42‰ to −37.21‰, including internationally recognized standards such as NBS19 and Carrara Marble.

### Taxonomic profiling from assembled contigs

Taxonomic composition was inferred from assembled contigs using CAT (Contig Annotation Tool)[7] 5.10.3. CAT assigns lineages to contigs using protein-level homology searches and a lowest common ancestor framework. Genus-level assignments were retained when support exceeded 0.70. Contig lengths were extracted from assembly FASTA files and aggregated by genus within each metagenome. Taxonomic profiles were calculated as the proportion of classified assembled base pairs assigned to each genus per metagenome. Taxa present in fewer than 20% of samples were removed prior to analysis to reduce sparsity. Because assembly recovery and classification success can vary with sequencing effort, sequencing depth (log-transformed total reads) was included as a covariate and controlled in downstream multivariate analysis. Profiles were Hellinger-transformed and summarized using principal component analysis (PCA); the first ten principal components (cumulative variance explained = 44.7%) were retained for downstream analyses. Variation partitioning (RDA; adjusted $R^2$) was used to quantify independent and shared contributions of environmental variables and taxonomic composition to GCF distributions while conditioning on sequencing depth.
